# RNA Binding of GAPDH Controls Transcript Stability and Protein Translation in Acute Myeloid Leukemia

**DOI:** 10.1101/2024.12.02.626357

**Authors:** Sama Shamloo, Jeffrey L. Schloßhauer, Shashank Tiwari, Kim Denise Fischer, Yohana Ghebrechristos, Lisa Kratzenberg, Aathma Merin Bejoy, Ioannis Aifantis, Eric Wang, Jochen Imig

## Abstract

Dysregulation of RNA binding proteins (RBPs) is a hallmark in cancerous cells. In acute myeloid leukemia (AML) RBPs are key regulators of tumor proliferation. While classical RBPs have defined RNA binding domains, RNA recognition and function in AML by non-canonical RBPs (ncRBPs) remain unclear. Given the inherent complexity of targeting AML broadly, our goal was to uncover potential ncRBP candidates critical for AML survival using a CRISPR/Cas-based screening. We identified the glycolytic enzyme glyceraldehyde-3-phosphate dehydrogenase (GAPDH) as a pro-proliferative factor in AML cells. Based on cross-linking and immunoprecipitation (CLIP), we are defining the global targetome, detecting novel RNA targets mainly located within 5’UTRs, including GAPDH, RPL13a, and PKM. The knockdown of GAPDH unveiled genetic pathways related to ribosome biogenesis, translation initiation, and regulation. Moreover, we demonstrated a stabilizing effect through GAPDH binding to target transcripts including its own mRNA. The present findings provide new insights on the RNA functions and characteristics of GAPDH in AML.

## Introduction

Acute myeloid leukemia (AML) is characterized by the uncontrolled proliferation of hematopoietic stem and progenitor cells, leading to neoplastic growth (1,2). The development of AML can arise from diverse mutations, including signaling and kinase pathway mutations and mutated epigenetic modifiers, directly influencing prognosis (3,4). Typically, a reported 5-year survival rate of only 30-35% underscores the urgent need for additional targeted therapies (5,6). As current therapeutic strategies predominantly target specific mutations, there’s an evident call for broader treatment approaches (7). In recent years, the prominence of dysregulated RNA-binding proteins (RBPs) in cancer, particularly their role in post-transcriptional regulation, has come to the forefront of cancer research (8–10). Given their involvement in critical processes such as protein translation, RNA splicing, RNA stability, and RNA localization, RBPs can drive cancer progression by disrupting cell proliferation and invasion mechanisms (11,12). Binding to RNA is often achieved with specific RNA binding domains (RBDs), including the RNA recognition motif, DEAD domain, or RGG/RG motif (13,14). However, many RBPs were identified lacking common RBDs, termed non-canonical RBPs (ncRBPs) (15–17). Previously, we identified several classical RBP candidates upregulated in AML, playing essential roles in RNA splicing and the survival of AML cells (18). Although numerous ncRBPs have been found, their roles in cancer physiology remain unclear (19). Notably, several metabolic enzymes engaged in glycolysis, the primary energy-generating pathway in cancer cells, also serve as ncRBPs such as pyruvate Kinase M (20), aldolase A (21), phosphoglycerate kinase (22) and GAPDH (23). The role of glycolytic enzymes is especially important in cancer metabolism, as during tumorigenesis there is an increased glucose uptake and reduced mitochondrial oxidative phosphorylation, known as the Warburg effect (24). The glycolytic enzyme glyceraldehyde-3-phosphate dehydrogenase (GAPDH) has been found to be upregulated in different cancer types, suggesting its role in tumor development (25–27). GAPDH is a known cancer vulnerability gene, and its depletion in various CRISPR-based viability screens has been observed in different cancer cell lines (28).

Beyond its pivotal role in glycolysis, GAPDH demonstrates a broad spectrum of functions, also referred to as its “moonlighting functions”, including membrane fusion activity, DNA repair and replication, modulation of iron metabolism, and regulation of cell death (29,30). Moreover, GAPDH was reported to bind RNA as a ncRBP, exhibiting a high preference for Adenine-Uridine-rich elements (AREs) on transcripts, such as the oncogene MYC and the inflammatory cytokine tumor necrosis factor-alpha (TNF-α) (23,31). It was hypothesized that GAPDH binds to its target RNAs via a tetrameric arrangement, utilizing its NAD^+^-binding region featuring a Rossmann fold or the positively charged groove formed by its monomer interface (23,32,33). Based on mutational analysis, a proposed model suggests that GAPDH binds to its target TNFα-ARE proximal to the Rossmann fold, at the dimer and tetramer interfaces via a sequential mechanism (34). However, the specific RNA-binding region remains a subject of ongoing research.

Within this study, we aimed to identify up-regulated ncRBPs permitting cellular survival and growth based on a clustered regularly interspaced short palindromic repeats (CRISPR)/Cas9 negative screen, targeting ncRBPs in AML cells. Considering the significance of GAPDH in AML, our objective was to pinpoint its RNA targets through CLIP and RNA-seq analysis following GAPDH knockdown. Additionally, we utilized biophysical assays to analyse the binding dynamics of GAPDH-RNA interactions and its consequential effect on RNA target transcript stability and translation.

## Materials and Methods

### Cell Lines and Cell Culture

Human MOLM-13 cells were purchased from DSMZ (Cat. No. ACC 554). The human THP-1 cell line was purchased from Abcam (ab275477). Cultivation was performed in RPMI 1640 medium containing 10% FBS and 1% penicillin/streptomycin at 37 °C and 5% CO_2_. Stable transfected cells contained 5 µg/ml puromycin to apply selective pressure. The Lenti-X™293T cell line from Takara (Cat. No. 632180) was utilized for lentiviral packaging.

### Lentivirus production and stable cell line generation

Knockdown and knockout of GAPDH was achieved by lentiviral transduction of shRNA and single guide RNA (sgRNA) into MOLM-13 cells and THP-1 cells, respectively. Lentivirus particles were produced by transient transfection into HEK293T cells as previously described (35). Briefly, 15 x 10^6^ HEK293T cells were seeded in a 15 cm plate a day prior to transfection to achieve 80–90% confluency the next day. 22.5 µg transfer vector, 11 µg VSVG vector (Addgene #8454), and 16.5 µg of pPAX2 vector (Addgene #12260) were transfected into HEK293T cells using 75 µg linear polyethylenimine (PEI, Polysciences). 48 hours post-transfection, the virus supernatant was collected, filtered with a 0.2 µm pore PTFE Membrane (Corning), and concentrated using centrifugal filter units. Concentrated viruses were used immediately or stored at -80°C. To produce stable shRNA producing cell lines the concentrated viruses were transduced into MOLM-13 cells and THP-1 cells followed by selection pressure with 5 µg/ml puromycin 48 hours post-transduction.

### CRISPR-Knockout screening

All sgRNA sequences were selected from the Brunello genome-wide library (4 sgRNAs/gene) (36). Individual sgRNA oligos were synthesized by Twist Bioscience on a 12K array and amplified using array primers (Array_Fwd: TTATATATCTTGTGGAAAGGACGAAACACCG; Array_Rev: AGCCTTATTTTAACTTGCTATTTCTAGCTCTAAAAC). Using a Gibson Assembly master mix (New England Biolabs), we cloned sgRNAs into a lentiviral sgRNA GFP-tagged vector (LRG) (Addgene #65656). Gibson reactions were transformed using DH10B electrocompetent cells (Invitrogen) at 2 kV, 200 Ω, and 25 uF. Bacterial colonies were quantified to obtain ∼70X coverage. Subsequently, library was deep-sequenced using MiSeq to confirm sgRNA representation. Cas9 cell lines were generated by transducing the cell lines, MOLM-13, THP-1 Jurkat and CUTLL1 with a lentiviral Cas9 construct and blasticidin resistance marker (Addgene #52962). Cell lines were transduced with the RBP library at a low MOI (∼0.3). On Day 4 post-transduction, GFP percentage was assessed to determine infection efficiency and sgRNA coverage (∼300-500X). Remaining 300-500X cells were placed back into culture after each passage until 20 days’ post-transduction. Genomic DNA (gDNA) of cells containing 300-500X coverage were harvested on Day 4 and Day 20 using Qiagen DNA kit based on manufacturer’s protocol. For library construction, 200 ng of gDNA was amplified for 20 cycles using Phusion Master Mix. End repair products were generated using T4 DNA polymerase (NEB), DNA polymerase I (NEB) and T4 polynucleotide kinase (NEB). Subsequently, A-tail overhangs were added using Klenow DNA Pol Exo-(NEB). DNA fragments were then ligated with Illumina-compatible barcodes (Bioo Scientific) using T4 Quick Ligase (NEB) and amplified using pre-capture primers (5 cycles). Barcoded libraries were then sequenced using MiSeq instrument (150 cycles). For pooled CRISPR screen analysis, individual time points for all samples were normalized using the formula (sgRNA read count/total read count) x 100,000. Subsequently, normalized reads were then used to calculate log2 fold change (normalized read count Day 4/normalized read count Day 20).

### AML-TCGA analysis

Raw RNA-seq reads from healthy control cells were acquired from the specified sources listed in the Supplementary Table S1. The TCGA pipeline method was utilized to quantify gene expression. Therefore, reads were aligned with STAR version 2.7.5 (37) utilizing the GRCh38 human assembly (38). The gene expression data of 151 AML patients from the TCGA project (produced by STAR concurrent with alignment) was obtained from GDC TCGA-LAML data portal (39). Healthy samples and TCGA raw counts were merged together. The R-package RUVSeq (40) was employed for batch correction, accounting for technical artifacts, before conducting the subsequent analysis of differential gene expression using the DESeq2 package (41). Differentially expressed genes were defined with a log fold change > 1 and multiple testing corrected p-value <0.1 (Benjamini-Hochberg). The PCATools (42) with the R package version 2.12.0 was utilized to validate the batch correction procedure by performing principal component analysis. EnhancedVolcano (43) with the R package version 1.18.0 was used to create volcano plots and batch corrected-vst transformed counts (from DESeq2) were used for the violin plot. The Wilcoxon’s rank-sum test was performed to test for statistical significance using a total of 151 tumor samples and 74 healthy control samples. Moreover, TCGA tumor data was divided into two subsets based on the expression level of GAPDH (expression ≥ median or expression < median) in VST transformed expression data (from DESeq2). Gene set enrichment analysis (44,45) with the R package version 1.20.3 was performed on all the TCGA-LAML samples and samples with high GAPDH expression using the Hallmark gene set (from msigdbr (46) with the R package version 7.5.1) along with the set of genes previously identified as binding targets of GAPDH from the eCLIP-Seq consensus peak. A correlation matrix was generated with the Spearman correlation of the expression of GAPDH with other genes. This matrix was used as the input for the gene set enrichment analysis.

### Knockdown and Knockout of GAPDH in AML cells

We established an inducible knockdown of GAPDH to assess the impact of its controlled depletion under optimal growth conditions. Knockdown of GAPDH in MOLM-13 cells was achieved by lentiviral transduction, as described above. Therefore, the transfer vector Tet-pLKO-puro (Addgene #21915) constitutively expressing the tetracycline repressor and puromycin N-acetyl-transferase, while the desired shRNA (shRNA 1 and shRNA 2) is driven by a Dox-inducible promoter, was utilized. Alternatively, GAPDH knockout cells were obtained by using the target vector TLCV2 (Addgene #87360) containing a Dox-inducible Cas9 enzyme fused to a GFP, a constitutively expressed reverse tetracycline-controlled transactivator fused to puromycin N-acetyl-transferase, as well an U6 promoter driving sgRNA (sgRNA 1-4, Table S2) expression to target GAPDH.

### GFP Competition Assay

The competition between parental MOLM-13, THP-1 and Jurkat cells and cells lacking GAPDH, RPS10, RPL15, MRPL32 or RPL18A, was monitored by a GFP competition assay. Therefore, 1.5 x 10^5^ MOLM-13 cells were infected with lentiviruses. The virus was produced by using the TLCV2 vector containing a Dox-inducible Cas9 enzyme fused to a GFP and a constitutively best performing sgRNA from the CRISPR targeting GAPDH, RPS10, RPL15, MRPL32 or RPL18A, as mentioned above. Stable transfected cells were induced with 5 µg/ml Dox. After two days, cells were washed twice with PBS and one third of cells were analyzed for GFP expression using the flow cytometry instrument SH800S (Sony), while two third of the cells were continuously grown over time. The growth medium was changed every two to three days to maintain Dox treatment. Measurement of GFP expression was performed after two, six, ten, fourteen, and seventeen after the first Dox treatment.

### Western Blot Analysis

Cells were washed with PBS prior to cell lysis with RIPA buffer (Thermo Fisher Scientific). The protein concentration was determined using the Pierce BCA Protein Assay Kit (Thermo Fisher Scientific) according to the manufacturer’s instructions. Proteins were separated using the SDS-PAGE system (Biorad) by loading 10 µg protein per sample. Following gel electrophoresis proteins were transferred to an Immobilon-P PVDF membrane (Merck Millipore) using the Wet/Tank Blotting System (Biorad). GAPDH was detected using an anti-GAPDH mouse monoclonal antibody from Sigma Aldrich (Cat. No. G8795) and a secondary anti-mouse HRP antibody from Sigma Aldrich (Cat. No. A9044). The anti-β-actin rabbit antibody from Abcam (ab8227) was a gift from Herbert Waldmann (Max Planck Institute of Molecular Physiology, Dortmund) and was utilized as internal control.

### Cross-Linking Immunoprecipitation-High-Throughput Sequencing

CLIP was performed according to Van Nostrand *et al*. (47). Briefly, MOLM-13 cells were UV crosslinked using one time 100 mJoule/cm^2^ at 254 nm radiation and two times 150 mJoule/cm^2^ at 365 nm radiation. Cell lysis was conducted using NP40 lysis buffer (50 mM HEPES, pH 7.5, 150 mM KCl, 0.5 % NP-40, 0.5 mM DTT, 2 mM EDTA) for 15 minutes on ice. Immunoprecipitation with an anti-GAPDH polyclonal antibody from proteintech (#10494-1-AP) was performed for one hour at 4°C while rotating. Cross-linked RNPs were T1 RNAse digested, RNA adapter ligated and blotted onto nitrocellulose membranes and monitored using a Typhoon Phospho-Imaging system (Amersham). One percent of the lysate input was saved and GAPDH protein detected by Western-blot as reference for GAPDH bound RNA quantification. Proteinase K treatment released RNPs were limited PCR amplified for Illumina sequencing with respective primer sets (47). Different metabolic conditions were analyzed using culture medium containing 11 mM glucose (norm), 5 mM glucose (low) or 11 mM galactose for 24 hours before cell lysis.

### RNA Extraction, Library Preparation and RNA-Sequencing

Total RNA was isolated from 3-5 × 10^6^ cultured MOLM-13 cells using TRIzol reagent (Thermo Fisher Scientific) according to the manufacturer’s instructions. For library preparation, we removed rRNA using the FastSelect-rRNA HMR Kit (QIAseq) following strand-specific cDNA library preparation with the Stranded Total RNA Lib Kit (QIAseq), in accordance to the manufacturer’s protocol. RNA sequencing was performed on Illumina NovaSeq S4 PE150 with at least 30 million reads yield per sample in the paired-end method. The zarp pipeline (48–51) was followed to process fastq files. FastQC (52), zpca (53) and MultiQC (54) were used for quality control. Adapters were trimmed through Cutadapt (55). The reads were mapped to the human genome (hg38, Genome Reference Consortium GRCh38) using STAR (37). Quantification was realized by using Salmon (56). The subsequent output was a count matrix, which was used as input for the R package DESeq2 (41) to identify differentially expressed genes. Genes with a log fold change >1 and adjusted p-value <0.05 were used for further analysis. The EnhancedVolcano R-package (57) with the R package version 1.18.0 was used to create the volcano plot. The ComplexHeatmap package (58) was utilized for creating heatmaps. The gene sets were obtained using the msigdbr (46) with the R-package version 7.5.1. The tools clusterProfiler (44) and enrichplot (45) with the R package version 1.20.3 were used for generating Gene Ontology and gene-set enrichment plots. PureCLIP (59) was performed to call crosslink sites. PureCLIP applies a hidden Markov model-based approach, which simultaneously performs peak-calling and individual crosslink site detection accounting for mutational profiling expected to occur at the crosslink sites. It incorporates a non-specific background signal and non-specific sequence biases. The peaks were generated by merging crosslink sites which in each other’s proximity and extending the solo crosslink sites up to 50bp for the downstream analysis. Exact mapping and peak calling parameters used are deposited under zenodo and can be found in the data availability section.

Cumulative distribution plots contain data from CLIP and knockdown of GAPDH, followed by RNA sequencing. The names of the genes were derived from the GAPDH clip peaks observed under various metabolic conditions. The empirical cumulative distribution function (eCDF) of the log fold changes for these gene sets was plotted alongside the distribution of log fold changes for all genes when comparing control versus knockdown across all metabolic conditions. Additionally, a negative control was included by randomly selecting genes from the list of all genes that did not code for RNA, which could be bound by GAPDH. The log fold change of GAPDH target genes was compared with the set of all genes using a Wilcoxon’s Rank sum test, and the fold change was calculated by using mean log fold change under different metabolic conditions. The final plot indicates these values (median, fold change, p-value). The PCA plots indicated that the variance introduced due to different metabolic conditions, and the samples were clustered together based on the knockdown of GAPDH. This was further validated when the number of differentially expressed genes in the condition specific comparisons was significantly low. Therefore, for the downstream analysis, the samples from different metabolic conditions were grouped together based on the knockdown of GAPDH. Adjusting the multiple testing correction using IHW.

### Recombinant Production and Purification of GAPDH

GAPDH (P04406, G3P_HUMAN) containing a N-terminal His-tag was expressed in BL21-CodonPlus (DE3)-RIPL *E. coli* cells (Agilent) using a pET-30 expression vector. Purification was performed by Ni-NTA agarose and size exclusion chromatography.

### Quantitative PCR

RNA was extracted from MOLM-13 cells using TRIzol (Thermo Fisher Scientific) according to the manufacturer’s instructions. cDNA was produced using the High-Capacity cDNA Reverse Transcription Kit (Applied Biosystems) and quantitative PCR (qPCR) performed with GoTaq qPCR Master Mix (Promega). The mRNA expression was detected using specific primer pairs as listed in Supplementary Table S2. mRNA levels were expressed relative to 18S rRNA. Relative gene expression was calculated using the 2^-ddCT^ method (60).

### Analysis of actively translated mRNAs

The expression of shLacZ and shGAPDH in MOLM-13 cells was induced with 2 µg/mL Dox. Total RNA was isolated using TRIzol (Thermo Fisher Scientific) according to the manufacturer’s instructions. The ribosome nascent-chain complex-bound mRNAs (RNC-mRNAs), also termed actively translated mRNAs, were isolated as described previously (61). Briefly, 5 x 10^6^ cells were incubated with 100 µg/ml cycloheximide (CHX) in culture medium at 37°C for 15 min. Afterwards, cells were washed two times using ice-cold PBS (containing 100 µg/ml CHX). Cell lysis was performed for 30 min in lysis buffer (20 mM Tris-HCl (pH 7.4), 10 mM MgCl_2_, 150 mM KCl, 100 μg/ml CHX, 1% Triton X-100 and 2 mM dithiothreitol) on ice. Cell debris was removed by centrifuging at 16.000 x g for 10 min at 4°C. The supernatant was carefully transferred on the surface of 20 ml sucrose buffer (30% sucrose in 20 mM Tris-HCl (pH 7.4), 10 mM MgCl_2_, 150 mM KCl, 100 μg/ml CHX and 2 mM dithiothreitol). Ultracentrifugation was carried out at 185.000 x g at 4°C for 5 hours. The supernatant was discarded and the pellet containing the RNC-mRNA was solubilized in Trizol (Thermo Fisher Scientific). RNAs for qPCR were treated as described above. mRNA levels were expressed relative to 18S rRNA. RNA derived from RNC-mRNA was compared to total RNA for both shLacZ and shGAPDH samples using the 2^-ddCT^ method (60). Moreover, RNAseq was performed as described above. The raw paired-end FASTQ files were first processed with Trimmomatic to remove low-quality bases and adapter sequences, improving overall data quality. Next, SortMeRNA was used to filter out rRNA reads, retaining only mRNA reads for a more accurate transcriptomic analysis. The resulting reads were then aligned to the human reference genome using STAR, a splice-aware aligner optimized for high-throughput RNA-seq data. Aligned reads were assigned to genes with featureCounts, generating a raw count matrix for downstream differential expression analysis. To examine translation efficiency (TE), we compared eCLIP target genes and control genes in GAPDH knockdown and LacZ control samples, with a focus on GAPDH, PKM, and RPL13. Additionally, TE across all genes was analyzed between the GAPDH knockdown and LacZ control conditions. Both analyses used the edgeR package for count normalization, generalized linear modeling, and statistical testing to identify differential TE.

### Fluorescence Polarization Assay

For the binding assay, increasing concentrations of GAPDH protein as a 2-fold dilution series in binding buffer (10 mM Tris-HCl, pH 8.0, 50 mM KCl, 2 mM DTT, 0.5 mM EDTA, 0.1 mg/ml bovine serum albumin, 0.1 mg/ml heparin) were titrated against Cy3-labeled RNA oligos. RNA derived from TNFα AU-rich element (TNFα-ARE_38_) or control RNA from rabbit β-globin Rβ_31_, purchased from Integrated DNA Technologies (Supplementary Table S2), were used at a concentration of 2 nM in a final volume of 20 µL in a black 384-well microplate (Corning® 4514). Reaction mixture was incubated at room temperature for 30 minutes and the fluorescence polarization (mP) was measured using a Tecan Spark® plate reader with excitation 535nm and emission 590 nm for Cy3. K_D_ values were calculated using GraphPad Prism 9 by fitting mP values and the corresponding GAPDH concentrations into a nonlinear regression model.

For the competition assay, a solution of GAPDH protein at a concentration of 500 nM and Cy3-labeled RNA at a concentration of 4 nM in binding buffer was incubated at room temperature for 30 minutes in the dark. The mixture was added to a 2-fold dilution series of the inhibitor to obtain a final concentration of 250 nM and 2 nM of the GAPDH protein and RNA, respectively. The measurement was performed after another 30 minutes of incubation. mP values were plotted against the corresponding inhibitor concentrations. All experiments were performed in triplicates.

### Electrophoretic Mobility Shift Assay (EMSA)

Complex formation between GAPDH protein and different target RNAs was investigated by EMSA. Recombinant GAPDH protein was produced at the Dortmund Protein Facility (DPF). Cy3-labeled TNFα-ARE and control Cy3-labeled rabbit β-globin (Rβ31) or scramble RNA were synthesized by Integrated DNA Technologies (IDT). Alternatively, the 5’UTR of GAPDH, and RPL13a were *in vitro* transcribed using T7 RNA Polymerase (Thermo Fisher Scientific) and the RNA was fluorescently labeled using the Fluorescein RNA Labeling Mix (Roche) according to the manufacturer’s instructions. The templates for *in vitro* transcription can be found in Supplementary Table S3.

For EMSA, recombinant GAPDH protein with increasing range of concentration was titrated into a constant concentration of a labeled RNA in binding buffer (10 mM Tris-HCl pH 8, 50 mM KCl, 2 mM DTT, 0.5 mM EDTA, 0.1 mg/ml Heparin, 0.1 mg/ml BSA, 10% glycerol, 40 U/ml RiboLock RNase Inhibitor (Thermo Fisher Scientific)) for 20 min at room temperature. Afterwards samples were run on a native 6% polyacrylamide gel for 120 min at 4°C and 200 V. The fluorescently labeled RNA was detected using the ChemiDoc MP Imaging System (Biorad).

### Mass Photometry

Biophysical characterization of the interaction of recombinant GAPDH protein and TNFα-ARE, as well as Rβ_31_ control RNA was achieved by measuring the particle size of assembled complexes. Binding reactions were performed in molar ratios of 100 nM GAPDH to 100 nM RNA and 100 nM GAPDH and 1 µM RNA prior to analysis. Therefore, the reaction was set up in binding buffer (10 mM Tris-HCl (pH 8.0), 50 mM KCl, 2 mM DTT, 0.5 mM EDTA) for 25 min at room temperature. The molecular mass of the reaction was determined by the TwoMP mass photometer (Refeyn) according to the manufacturer’s instructions. Briefly, 18 µl buffer was added to the center of a gasket located on a clean coverslip. Samples were diluted 1:10 by adding 2 µl to the buffer in the gasket after setting the focus. A calibration mix of Bovine serum albumin and Thyroglobulin was utilized to convert the measured contrast to molecular mass. Data were analyzed using the DiscoverMP software (Refeyn).

### RNA-Stability Assay

The stability of transcripts was analyzed using an actinomycin D stability assay. Prior to the assay, 10^7^ stable MOLM-13 cells expressing shRNA against GAPDH RNA (shRNA1) or control shRNA (lacZ) were cultivated in the presence of 2 µg/mL Dox for two days to induce shRNA expression. Two days post-induction, 2 µg/ml actinomycin D or DMSO as control were added to the cells and during the next 8 hours, cells were harvested every one hour for RNA extraction, cDNA synthesis and qPCR.

### GAPDH-Activity Assay

GAPDH activity was measured using the GAPDH Activity Assay Kit (Sigma) according to the manufacturer’s instructions. This assay quantifies the enzymatic conversion of Glyceraldehyde-3-Phosphate (GAP) to 1, 3-Bisphosphate Glycerate (BPG), generating a colorimetric product detectable at 450 nm, directly proportional to the enzyme’s activity level. Briefly, recombinant GAPDH protein incubated with a target RNA or immunoprecipitated GAPDH from cell lysate was combined with GAPDH Assay Buffer to create 50 µL of sample for each reaction (well). Subsequently, 50 µL of each sample was dispensed into a 96-well plate, followed by the addition of 50 µL of the reaction mix (GAPDH assay buffer, GAPDH developer, and GAPDH substrate). Absorbance at 450 nm indicating NADH production was measured using a microplate reader at 37°C in the kinetic mode for 60 minutes. Additionally, NADH standards were prepared and analyzed following the provided instructions. GAPDH activity was calculated by comparing the sample absorbance values to the NADH standard curve.

## Results

### Identification of non-canonical RBPs in acute myeloid leukemia

Alterations of RBP expression are associated with cancer cell proliferation through regulating oncogenes and tumor-suppressors (8). Previously, we found that a network of up-regulated classical RBPs in AML was controlling AML survival (18). However, less is known about ncRBPs and their role in AML cell survival. Our strategy for identifying potential druggable targets in AML involved utilizing CRISPR/Cas9 technology to pinpoint essential ncRBPs driving AML progression. Extending our previous RBP screening in AML(18), we utilized the census of RBPs from Gerstberger *et al*. (14) to extract 1,045 gene candidates not included in our initial CRISPR/Cas9 knockout screen. This selection aligns with other reports on *de novo* characterized RBPs (62–67,17), demonstrating an overlap of 24-62% in RBP hits (Supplementary Table S4). Among these 1,045 genes, 182 RBPs harbor a defined RBD, while the remaining RBPs lack a canonical RBD according to Gerstberger *et al* (14). We utilized a 41,000 sgRNA library with four sgRNA per RBP. This library was integrated into Cas9-expressing AML cell lines MOLM-13 and THP-1, as well as Cas9-expressing T-acute lymphoblastic leukemia (T-ALL) cell lines CUTLL1 via lentiviruses (Figure 1a).

**Figure 1:**
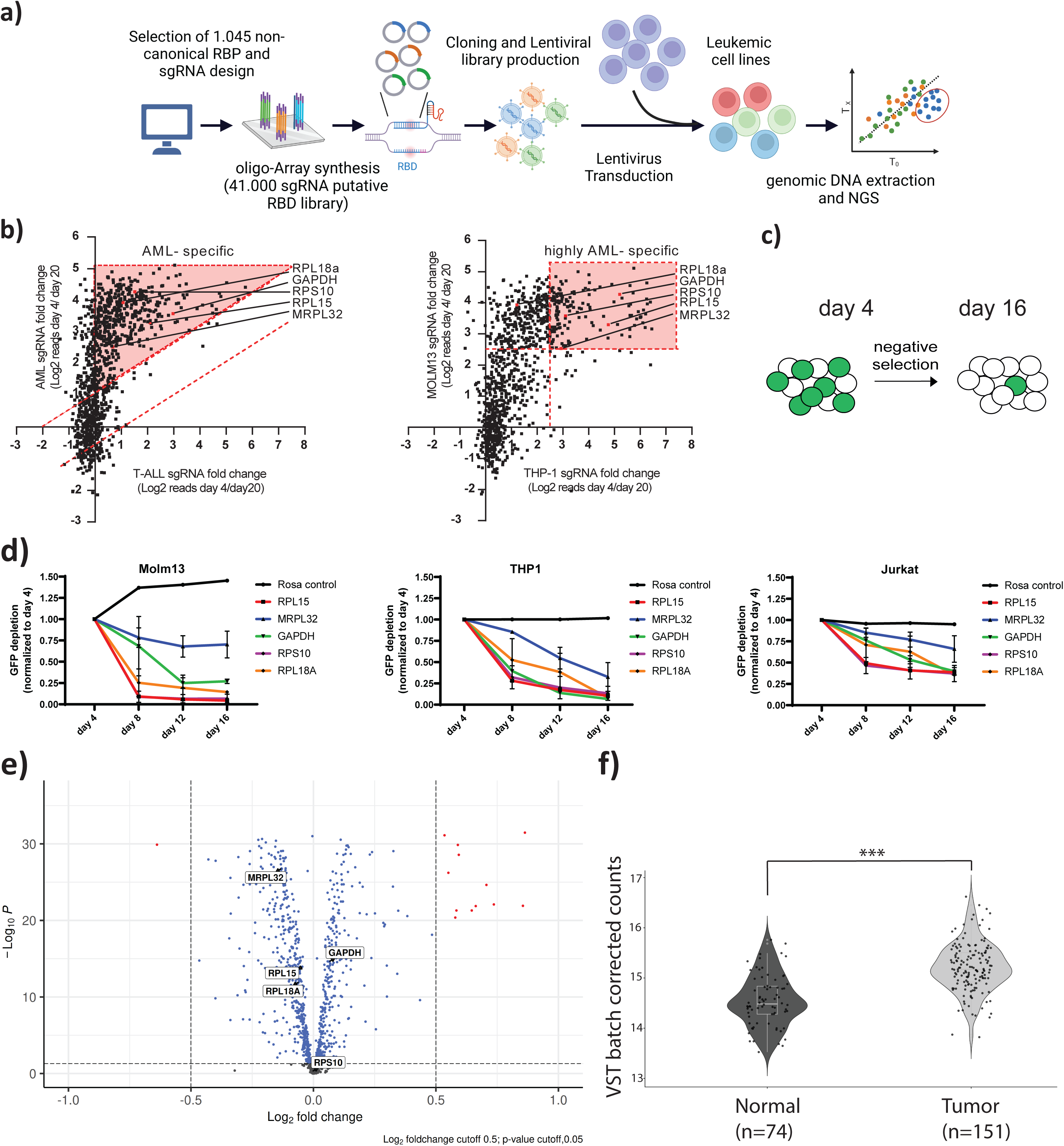
A CRISPR screen targeting non-canonical RNA binding proteins (ncRBPs) in leukemia cells. **(a)** Schematic presentation of the ncRBP-CRISPR screen. **(b)** Scatterplot of the ncRBP-CRISPR/Cas9 screening. The log2 fold change (fc) of sgRNA abundance (day 4/ day 20) for AML against T-ALL (left panel, dashed red lines and red area indicate selection threshold log2 fc > 1 each) and MOLM-13 cells against THP-1 cells (right panel, dashed red lines and red area indicate selection threshold log2 fc > 2.5 each) is plotted. Dots represent the average of sgRNAs targeting a ncRBP. Red dots indicate weak core fitness RBP genes. **(c)** Scheme of the GFP competition assay. **(d)** The AML cells (MOLM-13 and THP1) and T-ALL cells (Jurkat) were transfected by a sgRNA non-targeting vector (Rosa control) and sgRNA targeting vectors targeting the non-canonical RNA binding proteins RPL15, MRPL32, GAPDH, RPS10 and RPL18A. GFP fluorescence was monitored over time and the fluorescence signal normalized to day 4. **(e)** Volcano plot of differentially expressed ncRBPs in AML patient samples (n=151) compared to normal human CD34+ hematopoietic stem and progenitor cells (n=74). Vertical dashed lines indicate log2 fc cutoff 0.5. Genes above the horizontal dashed line have a p-value < 0.05. Grey dots indicate no observed significance, while blue and red dots indicate significant results. **(f)** GAPDH normalized counts based on the TCGA data using AML patient samples compared to healthy cells.

A loss-of-function pooled CRISPR screen was executed through next-generation sequencing, utilizing genomic DNA extracted on both day 4 and day 20 post-transduction. Key candidates were identified through a refined filtering process, highlighting hits with at least a five-fold depletion. Among the pool of 475 depleted RBP screen candidates, we identified 101 genes, which were reported to be weak core fitness genes in human cancer cell lines according to Bayesian Analysis of Gene EssentiaLity (68,69), which reflect potential suitable therapeutic (drug) targets. Small-scale analysis of the transcriptomes from both AML patients (n = 11) and healthy CD34^+^ hematopoietic stem cell (HSC) samples (n = 2) revealed differential expression patterns, with 31 genes down-regulated and 280 genes up-regulated in AML (Supplementary Figure S1). Regarding the filtered candidates, our analysis unveiled 16 out of the 101 weak core fitness genes demonstrating notable upregulation in AML to be considered for deeper analysis. Respecting previously identified weak core fitness genes, five ncRBPs were observed as AML specific hits, indicated by a counter-screen of T-ALL versus AML (Figure 1b, left). In-depth visualization of MOLM-13 and THP-1 using a counter-screen indicated 4 of 5 highly AML-specific hits, including GAPDH, RPS10, RPL15, and MRPL32 (Figure 1b, right). Although these proteins are crucial components of the ribosomes or play significant roles in metabolism, they appear to be core fitness genes in AML cells compared to T-ALL cells, suggesting a potential therapeutic window for targeting them. A GFP competition assay was conducted to verify the significance of GAPDH, RPS10, RPL15, MRPL32, and RPL18A in AML cell growth. Therefore, a knockout of the five ncRBPs was performed with subsequent analysis of GFP depletion over time (Figure 1c). Since GFP was co-expressed with the utilized sgRNAs, GFP depletion was analyzed by flow cytometry. If the analyzed sgRNA is essential for cell viability a decrease in GFP percentage can be monitored over time. Indeed, GAPDH, RPS10, RPL15, and RPL18A showed a drastic GFP fluorescence signal reduction after 16 days in both, MOLM-13 cells and THP-1 cells in contrast to T-ALL Jurkat cells (Figure 1d and Supplementary Figure S2).

The expression data in our analysis for ncRBPs was limited in terms of sample number. Therefore, we mined the expression level of all the 1045 candidates using a larger patient cohort. We analyzed datasets from The Cancer Genome Atlas (TCGA) project and other sources. In accordance with the differential expression of ncRBPs in AML patient samples (n = 151) compared to normal human CD34^+^ hematopoietic stem cells (HSC) (n = 74), GAPDH was slightly but significantly up-regulated (Figure 1e). However, RPL15, MRPL32 and RPL18A were significantly down-regulated, while RPS10 was only slightly overexpressed. The overexpression of GAPDH in tumor samples is further highlighted by visualization of GAPDH normalized counts based on the TCGA data (Figure 1f). We therefore focused to study GAPDH in depth as any reports pointed out the significant role in cancer-related processes (25,70,71). To strengthen the validity of GAPDH knockout on AML cells, we performed the GFP competition assay (72) using four different sgRNAs targeting GAPDH in MOLM-13 cells. Only 4% GFP-expressing cells could be observed by knocking out GAPDH 16 days post-induction, contrary to 53% and 60% GFP expressing cells using a sgRNA non-targeting control and empty vector control, respectively. The reduced survival rate of GFP positive cells observed in control conditions may be linked to the deleterious effects imposed by the overexpression of the Cas9-GFP fusion protein on cellular physiology.

In summary, our data demonstrate that selected targets, particularly GAPDH, are required for AML survival. Accordingly, we aimed to elucidate the role of GAPDH as a ncRBP in AML cells.

### GAPDH binds as a non-canonical RNA binding protein to various RNA species

Cancer cells primarily generate their energy in the presence or absence of oxygen through glycolysis and subsequent production of lactate instead of supplying pyruvate to the citrate cycle, known as the Warburg effect. It has also been proposed that RNA binding to GAPDH partially depends on the Rossmann fold and can be competitively affected by elevated levels of NADH/NAD^+^ (73). This implies that variations in energy levels produced by different glycolysis substrates and concentrations may impact RNA binding. It was shown that GAPDH binding in T-cells to IFN-γ and IL-2 mRNA was increased in cells not relying on glycolysis (74). Therefore, as GAPDH represents a key player in glycolysis, we aimed to shift the energy production from glycolysis to oxidative phosphorylation (OXPHOS), by cultivating the cells under reduced glucose condition or upon exchange of glucose to galactose to facilitate the analysis of RNA binding properties (75–77). On the one hand, we achieved RNA binding UV-crosslinked to GAPDH protein visualized by ^32^P-RNA labelling and autoradiography detection (Supplementary Figure S3). On the other hand, only slightly increased binding of GAPDH to RNA was observed, when cultivating cells under low glucose or galactose conditions. Nevertheless, we conducted cross linking RNA immunoprecipitation (CLIP) across the different metabolic conditions: normal glucose, low glucose, and galactose. This approach aims to map the global GAPDH-RNA targetome in AML and uncover how GAPDH may regulate target RNAs post-transcriptionally, depending on the cell’s energy state as reported previously in a global fashion (74). We performed two replicates for each metabolic condition. The total number of annotated peaks is 1774 with 163 peaks present in all conditions, 216 peaks present in only two conditions and 846 peaks are only present in one condition, suggesting partial targeting diversity dependent on metabolic state (Figure 2a, Supplementary Table S5). However, motif discovery revealed that GAPDH binds to the same putative motifs in all conditions, although a strong motif enrichment was not observed. This may indicate that GAPDH does not bind to its targets strongly in a sequence-specific manner as previously considered, but similar to other non-canonical RBPs (17) (Supplementary Figure S4). With ∼50% of CLIP hits, a high proportion of GAPDH targets were sequences associated with protein-coding located at the 5’UTR, the coding region and the 3’UTR (Figure 2b), while ∼20% of targets accounted for long non-coding RNAs (lncRNAs). Our data suggest a global pattern of GAPDH targets at diverse genomic regions, with a preference of binding to the 5’UTR of target transcripts (Figure 2c). Similar findings could be observed in all metabolic conditions (Supplementary Figure S5-S7).

**Figure 2:**
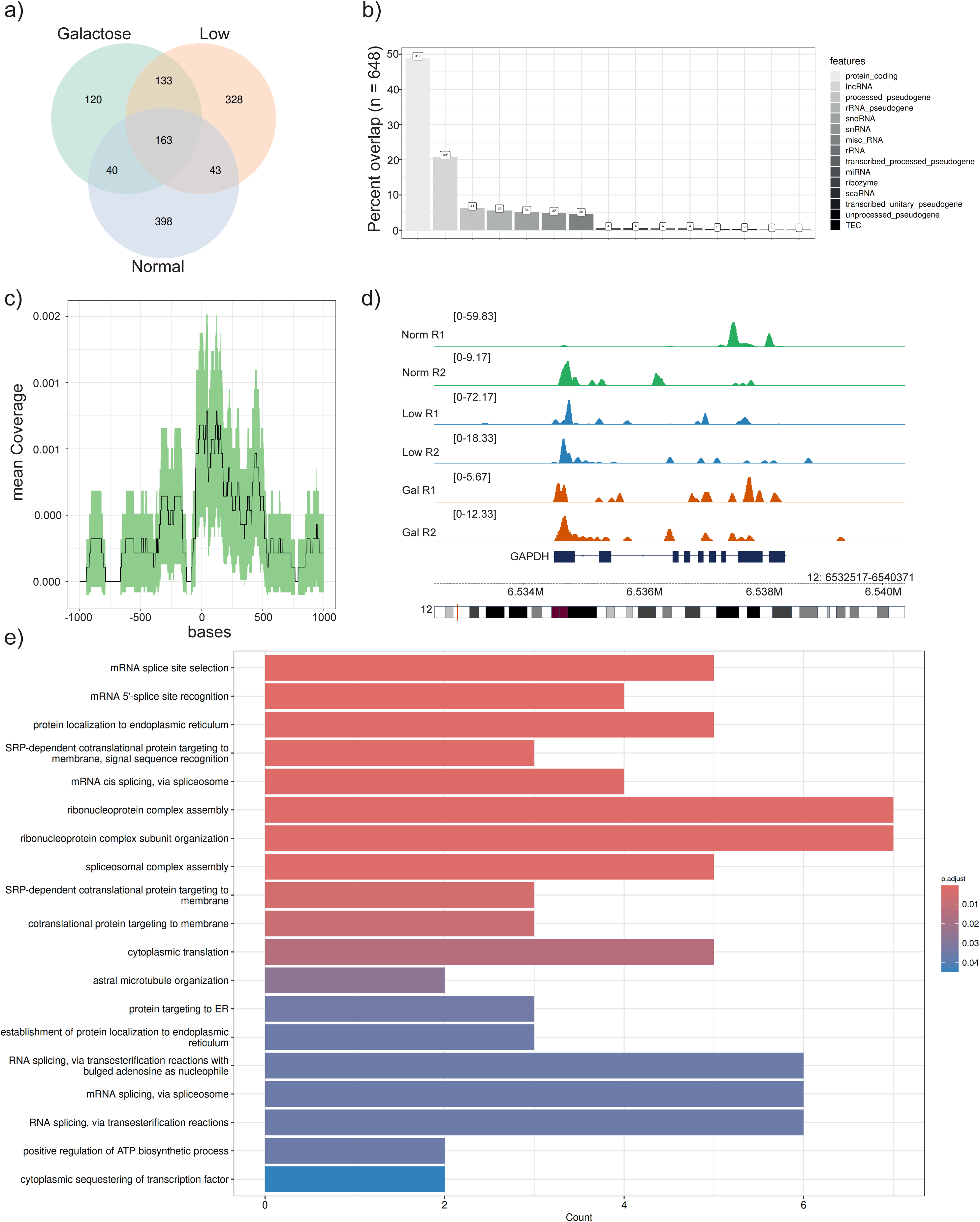
CLIP analysis of GAPDH binding sites in MOLM-13 cells. **(a)** Venn diagram of CLIP hits in galactose, low glucose and normal glucose conditions. **(b)** Genomic distribution of CLIP targets in normal glucose conditions. **(c)** Mean coverage of CLIP peaks in the 5’UTR in normal glucose conditions. **(d)** RNA-seq IGV plot of GAPDH and binding peaks in MOLM-13 cells in the presence of galactose, low glucose and normal glucose conditions. **(e)** Gene Ontology analysis based on CLIP. Gene Ontology analysis was conducted to identify common genes in all metabolic conditions.

We identified previously reported mRNA targets like TNFα (34) and MYC (23), though their target peaks did not align exactly with the previously described regions (Supplementary Table S6). Additionally, we discovered novel RNA targets, including the 5’UTR of GAPDH, the 5’UTR of the 60S ribosomal protein L13a (RPL13a), and Pyruvate Kinase M (PKM). However, their reported 3’UTR binding sites could not be completely recapitulated. Interestingly, GAPDH binds to its own transcript, indicated by the Integrative Genomics Viewer (IGV) tracks at different metabolic conditions (Figure 2d) and may have a self-regulatory function, a typical feature of many classical RBPs (78). Moreover, Gene Ontology analysis revealed involvement of GAPDH in important cellular functions such as protein localization, protein translation and splicing (Figure 2e). Gene Ontology analysis of unique and common to the different metabolic conditions can be found in Supplementary Figure S8-S11.

Next, we aimed to examine the effect of RNA binding on GAPDH activity to determine riboregulatory effects on its metabolic activity (79). Therefore, we immunoprecipitated GAPDH protein from MOLM-13 cell lysate and measured its enzymatic activity. However, no difference in GAPDH activity was observed between the RNase-treated and untreated samples (Supplementary Figure S12). Neither could we determine a riboregulatory effect using *in vitro*-synthesized RNA targets (Supplementary Figure S12) which is in contrast to other glycolytic enzymes (93).

### GAPDH is associated with the transcript stability and involved in protein translation

To assess the impact of GAPDH depletion overall and specifically on identified CLIP targets, we performed an induced GAPDH knockdown in AML cells using two distinct small hairpin RNAs (shRNAs). Induced knockdown of GAPDH led to decreased GAPDH mRNA and protein levels, while the control shRNA targeting LacZ had no effect (Figure 3a and 3b). The transcriptome was further analyzed upon shRNA knockdown of GAPDH at normal and low glucose levels, or in the presence of galactose instead of glucose in analogy to the CLIP assay. However, no major differential gene expression was observed under tested metabolic conditions under GAPDH knockdown. The PCA plots indicated that the variance introduced due to different metabolic conditions, and the samples were clustered together based on the knockdown of GAPDH (Supplementary Figure S13). This was further validated by the significantly low number of differentially expressed genes found in the condition-specific comparisons. We assume that the time of cultivating cells in the tested metabolic conditions may not be sufficient to observe a pronounced difference. For downstream analysis, samples from different metabolic conditions were grouped together based on the knockdown of GAPDH. RNA-seq analysis demonstrated the upregulation of 72 genes and downregulation of 104 genes, including GAPDH (Figure 3c). Using gene set enrichment analysis (GSEA) the GAPDH knockdown resulted in the activation of protein translation associated pathways including ribosomes biogenesis, initiation and regulation of translation (Figure 3d). Notably, cardiac-related gene sets were also significantly enriched (see also Supplementary Figure S14). In contrast, genes involved in nucleosome assembly and chromatin remodeling were significantly suppressed (Figure 3e). RNA-seq data following GAPDH depletion also indicate its involvement in protein translation. Importantly, it is noteworthy that the pathways are influenced by GAPDH’s diverse functions and cannot be solely attributed to its RNA-binding role. Consequently, we employed a cumulative distribution analysis to illustrate the variation in expression levels between the direct GAPDH CLIP targetome and a set of randomly selected non-target genes upon GAPDH knockdown in MOLM-13 cells (Figure 3f and Supplementary Figure S15). We intersected the identified binding sites from all CLIP experiments with the RNA-seq data after knockdown of GAPDH. A significant negative shift in the cumulative distribution function of the fold changes was evident across all tested metabolic conditions, though the *de novo* targets were distinct (Figure 3f and Supplementary Figure S15). The absence of GAPDH had a direct effect on the binding sites, indicating that the identified CLIP targets are real binding sites of GAPDH and GAPDH stabilizes its transcript targets as they are less abundant in the KD-RNA-Seq data. Similar findings could be determined for all individual metabolic conditions (Supplementary Figure S15).

**Figure 3:**
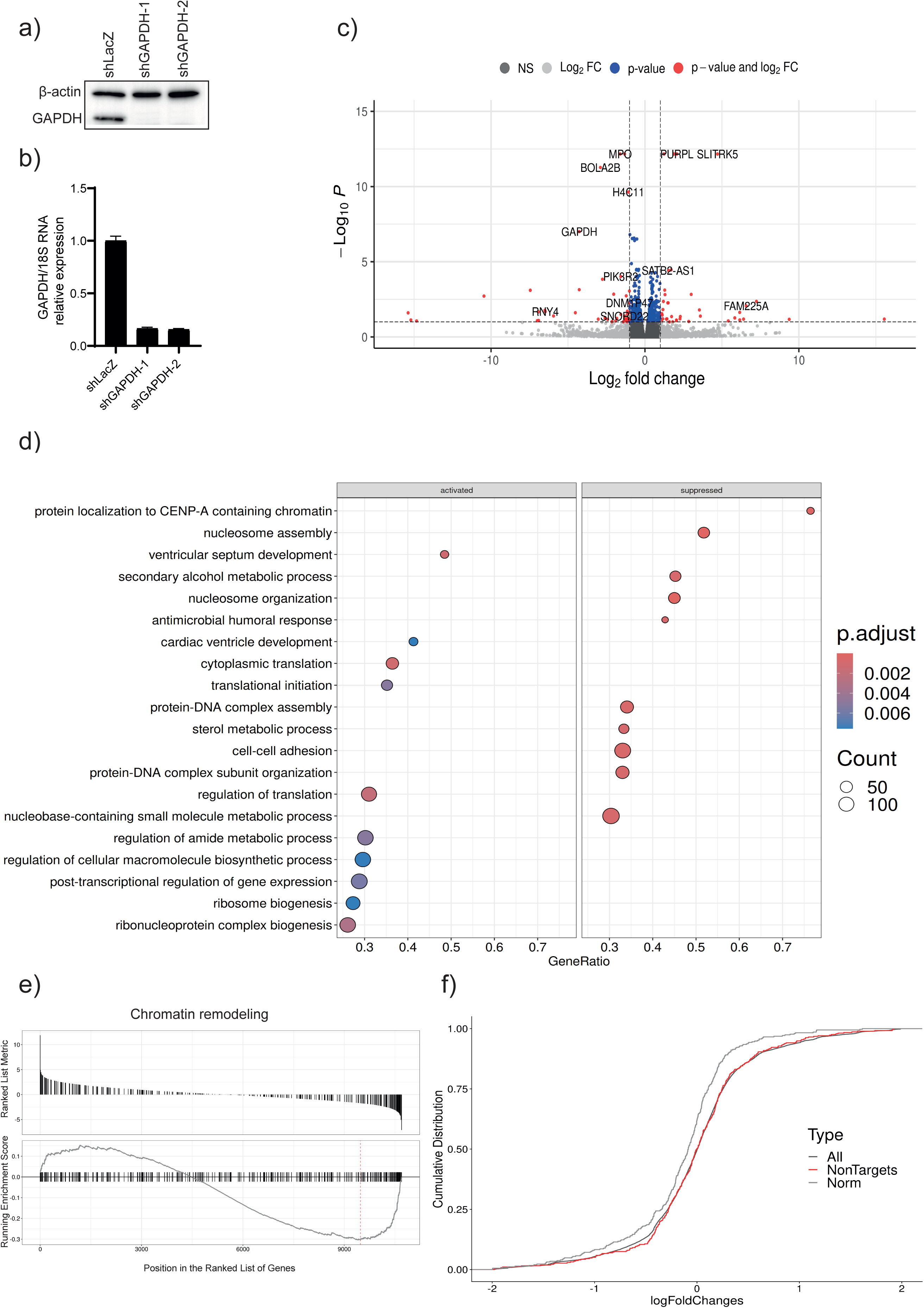
Knockdown of GAPDH in MOLM-13 cells. **(a)** Western blot of GAPDH in the presence or absence of Doxycycline (dox) using MOLM-13 cells targeted by shGAPDH-1, shGAPDH-2, and a shLacZ targeting control. β-actin was utilized as loading control. **(b)** Visualization of mRNA expression of GAPDH normalized to 18S rRNA after treating MOLM-13 cells with dox. **(c)** Volcano plot of GAPDH knockdown versus control MOLM-13 cells. Vertical dashed lines indicate log2 fold-change cutoff 1, p-value-cutoff 0.1. **(d)** Gene Ontology (GO) enrichment analysis of activated and suppressed pathways after GAPDH knockdown in MOLM-13 cells. **(e)** Enrichment plots from Gene Set Enrichment Analysis (GSEA) of chromatin remodeling genes. **(f)** Cumulative distribution function (CDF) plot for joined peaks under normal (Norm) glucose conditions. Bindings sites based on CLIP data were intersected with the RNA seq after GAPDH knockdown.

### Characterization of GAPDH binding to RNA targets

Based on the CLIP data, we intended to further validate the interaction of the identified GAPDH-RNA targets TNFα, GAPDH, and RPL13a with GAPDH protein. For this purpose, the electrophoretic mobility shift assay (EMSA) was utilized to assess the affinity of the interaction by varying the concentration of recombinantly expressed GAPDH in the presence of free RNA. Among the analyzed RNA targets, the AU-rich RNA derived from TNFα (TNFα-ARE) exhibited the highest affinity to GAPDH, with a K_D_ of 217 nM (Figure 4a). Further EMSA data including GAPDH 5’-UTR (K_D_= 0.4 µM) target site, RPL13 5’-UTR (K_D_ = 887 µM) and an additional scramble negative control can be found in Supplementary Figure S16 confirming our identified CLIP targets.

**Figure 4:**
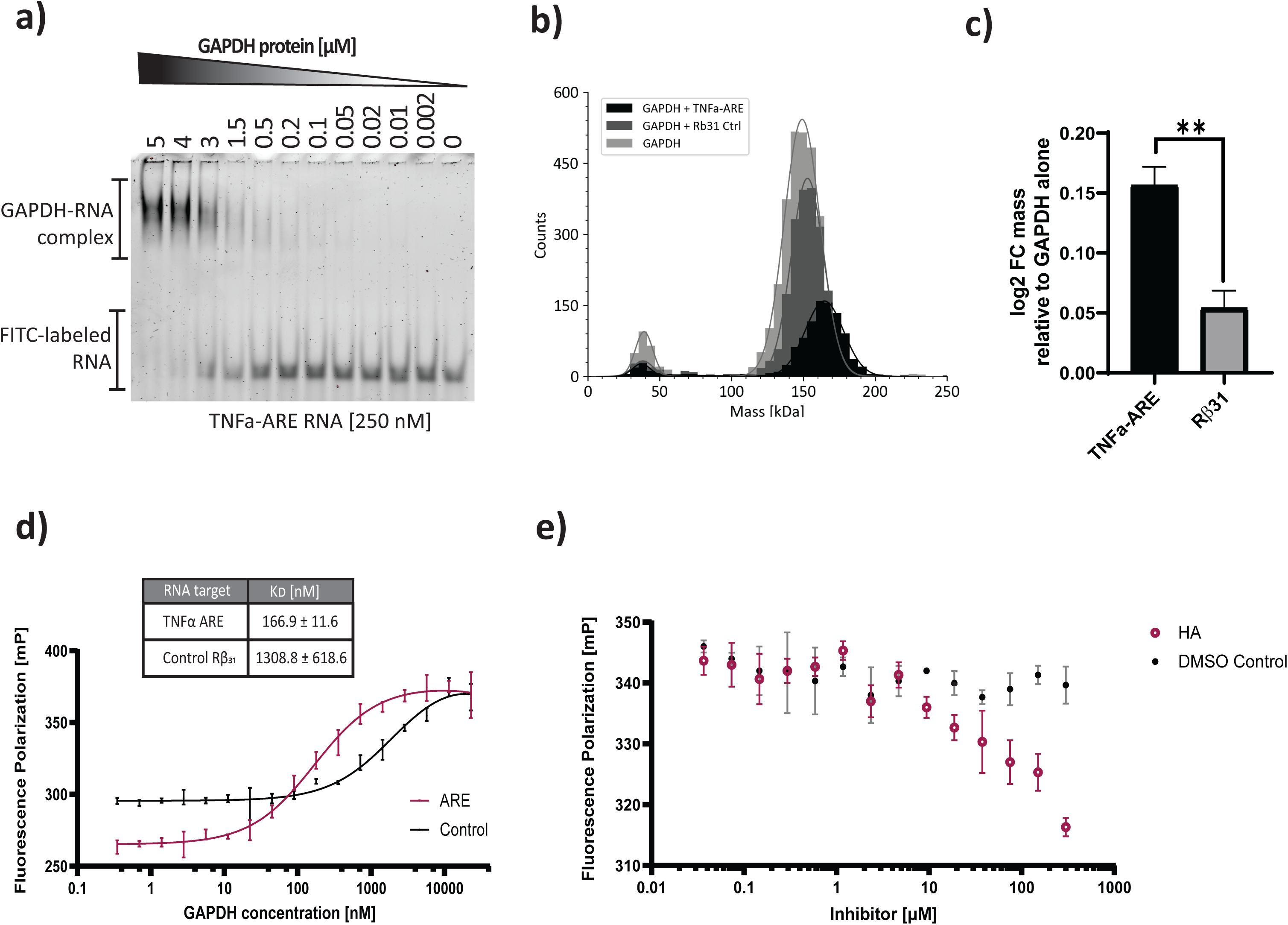
Characterization of GAPDH-RNA binding. **(a)** Electrophoresis mobility shift assay (EMSA) based on Cy3-labeled TNFα-ARE and GAPDH. **(b)** Overlay of mass photometry histograms generated from samples without RNA, in the presence of TNFα-ARE, or Rβ31 control. **(c)** Bar plot based on mass photometry data focusing on the tetramer peaks of TNFα-ARE or Rβ31 control relative to GAPDH alone. The data are expressed as log2 fold change (FC) mass. Data (n = 3) are shown as mean ± SD.** P < 0.01. **(d)** Fluorescence polarization assay of Cy3-labeled TNFα-ARE and Rβ31 control. **(e)** Fluorescence polarization assay of Cy3-labeled TNFα-ARE in the presence or absence of heptelidic acid (HA).

Our next objective was to elucidate the oligomeric state by which GAPDH binds to RNA through the utilization of mass photometry. Mass measurements revealed that the GAPDH monomer (38.5±0.4 kDa) and tetramer (148.3±0.3 kDa) exhibited mass values in line with the expected sizes of 37.5 kDa and 150 kDa, respectively. In the presence of TNFα-ARE, a noticeable shift of 17±0.3 kDa to ∼165-kDa total mass indicative of the formation of the GAPDH tetramer-RNA complex was evident (Figure 4b). Although a minor shift was noticeable when adding control rabbit β-globin 31 (Rβ31) to GAPDH, a significant increase of mass of the TNFα-ARE-GAPDH tetramer complex was observed (Figure 4c).

A fluorescence polarization assay was performed to elucidate the RNA binding behavior of GAPDH in the presence of the irreversible GAPDH inhibitor heptelidic acid (HA). First, the polarization assay was used to verify the findings of the EMSA. In contrast to the control RNA from Rβ31with a K_D_ of 1309 nM, TNFα-ARE demonstrated a notably higher affinity, with a K_D_ of 167 nM (Figure 4d). The observations are consistent with a previous study by White *et al*. (34), which yielded a K_D_ of 97 nM (TNFα-ARE) and 1400 nM (Rβ31). In the presence of HA, a concentration-dependent reduction of RNA binding could be observed, with a maximum of around 30% at 300 µM concentration (Figure 4e). We also examined different GAPDH targeting inhibitors 4-octyl itaconate (4-OI) and CGP3466B (Supplementary Figure S17) showing no inhibitory effect on RNA binding.

### GAPDH binding to RNA affects target RNA stability and target protein translation

The current data reveal binding of GAPDH to its own transcript. Consequently, we aimed to investigate GAPDH’s protein potential autoregulation on its mRNA stability. To accomplish this, we conducted an Actinomycin D RNA stability assay. Actinomycin D functions by intercalating with DNA within cells, thereby inhibiting the transcription of new mRNA (80). In the absence of Actinomycin D, the GAPDH knockdown mediated by shRNA led to a gradual reduction in GAPDH mRNA levels over time in contrast to control cells as expected (Figure 5a). Conversely, the administration of Actinomycin D to GAPDH knockdown cells elicited an exponential decline in GAPDH mRNA levels, while GAPDH mRNA was depleted linearly. GAPDH knockdown cells displayed a marked threefold reduction in the half-life of GAPDH transcripts compared to control cells (Figure 5b). These findings suggest a notable stabilizing effect of GAPDH on its own transcript. Moreover, GAPDH slightly stabilized the proto-oncogene MYC and TNFα transcripts (Supplementary Figure S18).

**Figure 5:**
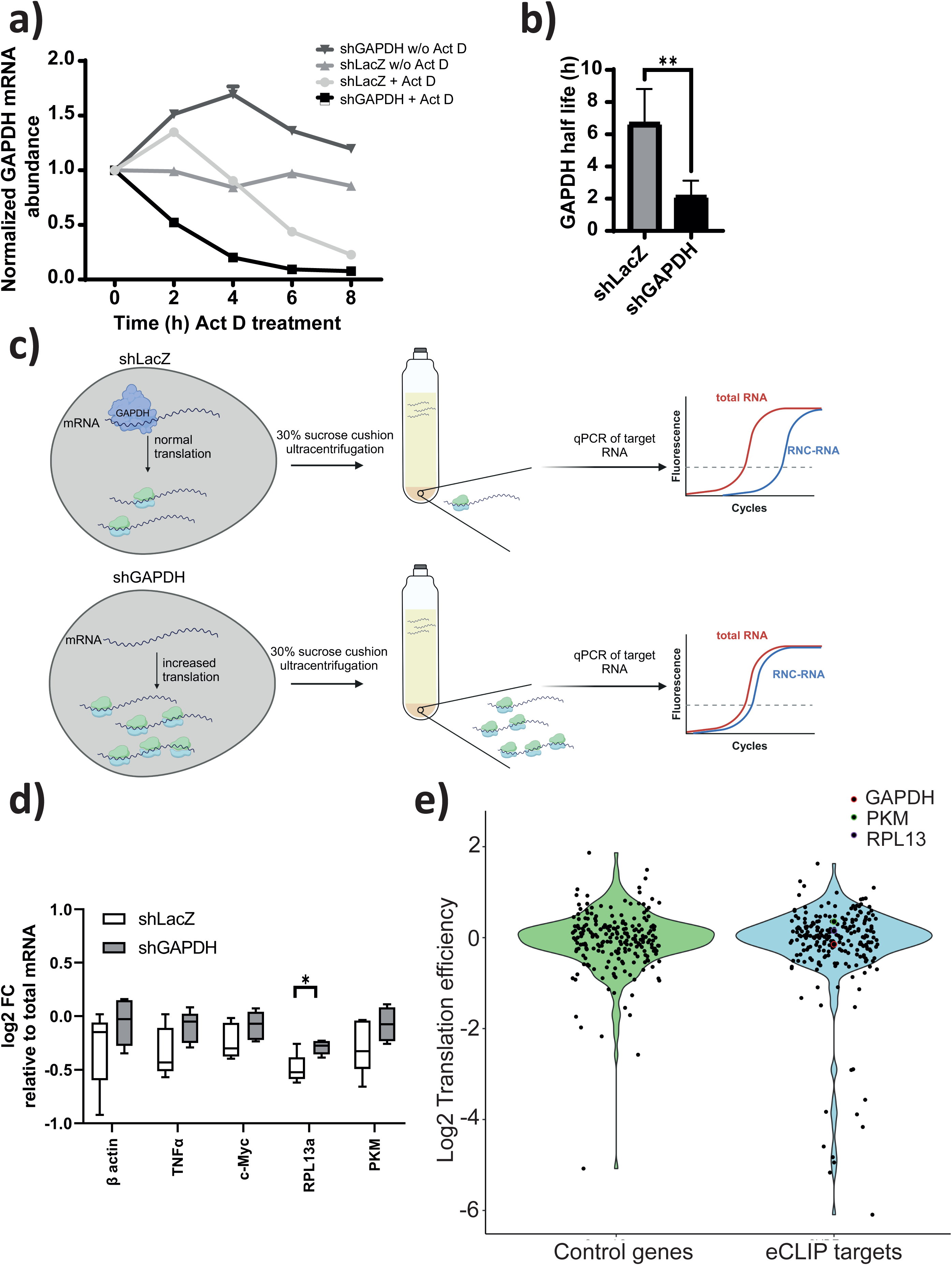
Characterization of GAPDH-RNA binding. **(a)** GAPDH mRNA expression of modified MOLM-13 cells utilizing an Actinomycin D (Act D) stability assay. MOLM-13 cells, expressing shLacZ and shGAPDH were analyzed in the presence (+) or absence (w/o) of ActD. **(b)** Bar blot visualizing GAPDH mRNA half-life based on the Act D assay. Data (n = 4) are shown as mean ± SD. **(c)** Scheme illustrating the isolation of actively translated mRNA. **(d)** Visualization of the relative mRNA expression levels of GAPDH targets in the nascent-chain complex-bound mRNA (RNC mRNA) fraction compared to the total mRNA. Samples were normalized to 18S rRNA and mRNA levels are expressed as log2 fold change (FC) using ddCt values with total RNA as calibrator. Data (n = 5) are shown as mean ± SD. * p < 0.05, ** p < 0.01. Created with BioRender.com. (e) Translational efficiency between eCLIP target genes and control genes based on RNAseq data. GAPDH (red), PKM (green), and RPL13 (purple) are highlighted.

We observed a prominent peak localization of GAPDH in the 5’UTRs of its target transcripts by CLIP (Figure 2c) and enriched GO terms of GAPDH knockdown in ribosome biogenesis and translational initiation (Figure 3d) which could imply a function in protein translation. To test this hypothesis, we elucidated GAPDH’s role in modulating protein translation of RNA targets, by analyzing the translation efficiency of GAPDH RNA targets in both the presence (shLacZ) and absence of GAPDH (shGAPDH). Therefore, MOLM-13 cells were treated with Dox to induce GAPDH knockdown. Afterwards, isolation of ribosome nascent-chain complex-bound mRNAs (RNC-mRNAs) according to Wang *et al*. (61) was achieved by ultracentrifugation (Figure 5c). The effect of GAPDH binding to RNA targets was analyzed by qPCR. The comparison of actively translated mRNA (RNC-mRNA) to total RNA corresponds to the translation efficiency. In the absence of GAPDH, the identified CLIP-targets TNFα, MYC, RPL13a, and PKM exhibited increased translation. It has to be emphasized, that the non-target β-actin also showed increased translation (Figure 5d), however being in line with the large scale sequencing analysis (Figure 5e). While only the difference for RPL13a is significant, a more precise in-depth analysis of CLIP targets was performed. Therefore, RNA-Seq of RNC-mRNA and total mRNA was conducted. The translation efficiency was calculated and eCLIP targets compared to a control gene set (Figure 5e). While GAPDH mRNA was reduced, PKM and RPL13a led to minor increased translation consistent with the qRT-PCR. It is noticeable, that a high number of eCLIP targets were either strongly positively or negatively reduced pointing to a very transcript and context dependent mechanisms on GAPDHs capacity to regulate translation possible involving diverse cis-acting factors. However, there was no absolute significant trend in either decrease or increased TE. A list of genes with TE value above +1 and below -1 can be found in the Supplementary Figure S19. Additionally, ribosome-associated genes exhibited reduced translation in the absence of GAPDH.

## Discussion

The dependence of cancer cells on GAPDH due to the increased glucose consumption is well known (81). The emerging roles of GAPDH in RNA binding within cancer cells underscore its critical involvement in promoting cancer cell proliferation, rendering it an attractive target for drug development. Yet, our understanding of the implications of GAPDH-RNA binding remains limited. Several studies have demonstrated GAPDH’s interaction with viral RNAs (82,83), tRNAs (84,33), ribozymes (85,86) and mRNAs (23,31,74). In the present study, we found a preferred binding of GAPDH to mRNAs, but also lncRNAs. Although GAPDH was reported primarily binding to ARE in the 3’UTR of some mRNA transcripts, as previously reviewed (29), we found a preference for binding in the 5’UTR on a global scale. The interaction of GAPDH with the 5’UTR may affect mRNA stability and the recruitment of the translation machinery, thereby changing protein production rates (87). Alteration of protein translation is crucial in cancer, empowering cancer cells to operate independently of growth factors and drive unregulated cell proliferation (88,89).

In line with this investigation, prior studies identified the pro-inflammatory TNFα-ARE as a target of GAPDH in various cell lines including AML cells and *in vitro* (34,90,31). Consistent with earlier research (23), the proto-oncogene MYC was recognized as a target of GAPDH, directly influencing the growth and proliferation of cancer cells (91). Contrary to prior findings indicating GAPDH binding primarily in the 3’UTR of TNFα and MYC, our research unveils its binding affinity within the 5’UTR. Interestingly, the 5’UTR of RPL13a mRNA was found as a GAPDH target. This finding is noteworthy, as GAPDH appears to play a dual role in regulating RPL13a, both at the mRNA level by targeting its 5’UTR and at the protein level by protecting it from proteasomal degradation (92).

In the present work, the mRNA of the glycolytic enzyme PKM was found to act as consistent GAPDH target. While other glycolytic enzymes, such as ENO1 has been shown to have a riboregulatory effect (93), our data indicate that GAPDH is not riboregulatory per se (Supplementary Figure S12). This highlights GAPDH’s moonlighting RNA-binding function in a one-way direction towards post-transcriptional gene regulation on the target transcripts and not vice versa acting on the energy metabolism. Whether additional metabolic enzymes evolved into diverse RBP functioning and if they can be classified in a similar fashion remains to be determined. Shen *et al*. correlated PKM and ENO1 genes with GAPDH expression in cancer cells (94). Moreover, the RNA binding abilities of ENO1 and PKM have been linked to cancer progression and proliferation (95,20,93). It is intriguing to speculate that other glycolytic enzymes fulfill a similar ncRBP function concerting together in glycolysis regulation (96,97). Further research is needed to evaluate the regulatory network of the glycolytic enzymes in AML.

Previous studies revealed either a stabilizing or destabilizing effect on GAPDH targets under various conditions (98–101). Here, we uncovered an auto-regulatory and auto-stabilizing role of GAPDH, supported by both CLIP/RNA-Seq analysis and the Actinomycin D assay. Stabilizing its own transcript while being slightly overexpressed may have positive implications for fueling glycolysis in AML and other cancer per se. Modulating specifically the stability and decay of GAPDH targets may be a promising cancer treatment strategy (102). However, more structural elucidation of the GAPDH RNP and its mode of function is necessary. One step forward in that direction was our attempt to test known GAPDH inhibitors for potential RNA binding interference. Only HA, a low-affine non-reversible GAPDH inhibitor, targeting the cysteine residue near the active site within the sequence of the S-loop domain of the enzyme, showed partial efficacy (103), which is also in the vicinity of the NAD^+^ binding site. We could demonstrate that modification of the cysteine residue led to decreased RNA binding, indicating that either the active site is at least partially involved in direct RNA binding or that HA confers an unfavorable confirmation change by communicating with the RNA binding interface including the NAD^+^ binding pocket (23). This finding might serve as a starting point to design a more selective GAPDH RBP inhibitors to dissect GAPDH’s functions in energy and RNA metabolism. Also, the other used GAPDH inhibitors targeting different protein site were ineffective to influence RNA binding unveiling that those protein domains are not required for RNP formation.

Consistently, our data, derived from direct analysis of actively translated CLIP targets following GAPDH knockdown, revealed an increased protein translation for few targets, including RPL13a and PKM. Taking a broader view of mRNA translation, RNA-seq data from translation efficiency assays revealed both upregulation and downregulation of translation across various targets, consistent with previous findings. Observations indicated that GAPDH represses protein translation by binding to the TNF-3’UTR (31), IFNγ-3’UTR (74) and the 3’UTR of the angiotensin II type 1 receptor (AT1R) (104).

Interestingly, GAPDH depletion had a negative effect on translation efficiency for ribosome-associated RNAs, highlighting it functions on general protein synthesis. Moreover, the RNA-seq analysis following GAPDH knockdown in MOLM-13 cells showed an upregulation of pathways involved in protein translation. Although, these findings may also be a result of a compensatory effect due to metabolic stress, GAPDH’s effect on translation was confirmed by the regulation of eCLIP targets.

Apparently, GAPDH acting as an RBP may have multiple functions on transcript stability and translation, depending on the cellular context, cis-acting factor and binding site preference presuming a highly complex network interaction with varying outcomes. Nevertheless, our data implies that GAPDH stabilizes its target transcripts by binding the 5’UTR in AML, potentially resulting in a modulation of ribosome association.

Furthermore, the reduction of GADPH led to the activation of cardiac related gene sets. This effect could be attributed to AT1R, which plays a significant role in hypertension and heart failure (104). Additionally, another study also elucidated a destabilizing effect of GAPDH on endothelial-derived vasoconstrictor endothelin-1 mRNA, which is a key player in vascular homeostasis (101). Here, we noticed that GAPDH knockdown resulted in the suppression of chromatin remodeling, demonstrating further nuclear moonlighting functions of GAPDH. Translocation of GAPDH into the nucleus and its regulatory function of gene expression was reported previously (107). Modification of the chromatin structure and genome instability was shown to be associated with GAPDH in yeast (108). Histone acetylation was promoted by GAPDH due to transnitrosylation of histone deacetylase, leading to chromatin remodeling (71,109). Direct binding of GAPDH to telomeres regulated chromosome stability, facilitating cancer cell proliferation (110,111).

We identified long lncRNAs as another large group of GAPDH RNA targets. However, the mechanisms underlying GAPDH’s interaction with lncRNAs and its physiological implications remain poorly understood. To date, our understanding of the mechanism by which GAPDH binds to its RNA targets is limited due to the lack of available GAPDH-RNA structures. In the future, studying GAPDH mutants that are deficient in RNA binding and clearly dissecting this function from its metabolic and other roles may enhance our comprehension of GAPDH’s as a ncRBP. However, this may not fully achievable due to the described overlap between partial RNA and NADH binding within the Rossmann fold (23). Nonetheless, our findings demonstrate the binding of TNFα-ARE by to fully catalytically active GAPDH tetramers, consistent with the observations made by White *et al*. (34).

In sum, here we successfully CRISPR screened ncRBPs associated with AML cell survival. We found the glycolytic enzyme GAPDH as a proliferative associated gene in AML cells and elucidated new binding sites and confirmed reported targets of GAPDH in AML cells. Moreover, pathways associated with GAPDH knockdown were unveiled. Finally, we could demonstrate GAPDH’s positive impact on transcript stability and impact regulation of translation of selected genes.

## Data Availability

The TCGA data underlying this article are available in the article and in its online supplementary material. For the CLIP-data the Gene Expression Omnibus (GEO) accession number is GSE270168 and the corresponding secure token is “qrizysqihjybpin”. For the RNAseq data after GAPDH knockdown the GEO accession number is GSE270169 and the corresponding secure token is “ydobyumwjtulzml”. The utilized source code for the eCLIP analysis can be found at https://zenodo.org/records/12170459?token=eyJhbGciOiJIUzUxMiJ9.eyJpZCI6IjdmYTBmMzk2LTBkNjktNDA0ZS04ZmJmLTk1ZDJjYTM3MmE0OCIsImRhdGEiOnt9LCJyYW5kb20iOiIwNjIwNDNlZDkzM2M5NTlkOTdhYjI0YWM0MmExYTY3MSJ9.fY0qGWo7nnjcwpM5F5zhi6-2kmbyZyqEyrb7BgJ_Kl-d0WyVvuOUh60rsjAlX1BNVWldRPJFfkMJoH_q7AXEig

## Acknowledgement

The authors thank Raphael Gasper-Schoenenbruecher (Max Planck Institute of Molecular Physiology, Dortmund) for his support in mass photometry. Moreover, special thanks go to Alicia Peschel (Max Planck Institute of Molecular Physiology, Dortmund) for her support with ultracentrifugation. The results here are in part based upon data generated by the TCGA Research Network: https://www.cancer.gov/tcga. We acknowledge A. Heguy and the NYU Genome Technology Center for expertise with sequencing experiments. We thank all past and present members of the Aifantis and Imig Labs for excellent scientific discussion, continuous support and intellectual contributions. We are grateful to Prof. Daniel Summer and Tzu-Chen Lin, TU Dortmund granting access and help to the flow-cytometry infrastructure. We thank Prof. Raunser, MPI Dortmund for access to BL-2 lab space.

## Author contributions

Conceptualization: JI, EW; Methodology: JI, EW, SS, ST and JLS; Investigation: JI, EW, SS, ST, JLS, KDF, YG, IA and LK; Formal analysis: JI, EW, SS, ST, JLS and KDF; Writing - Original Draft: JLS; Writing - Review & Editing: JLS, SS, and JI; Visualization: JI, EW, SS, ST, JLS; Project administration: JI; Funding acquisition: JI, AI

