## Supplemental Figures for "RNA Binding of GAPDH Controls Transcript Stability and Protein Translation in Acute Myeloid Leukemia"

### Supplementary Materials

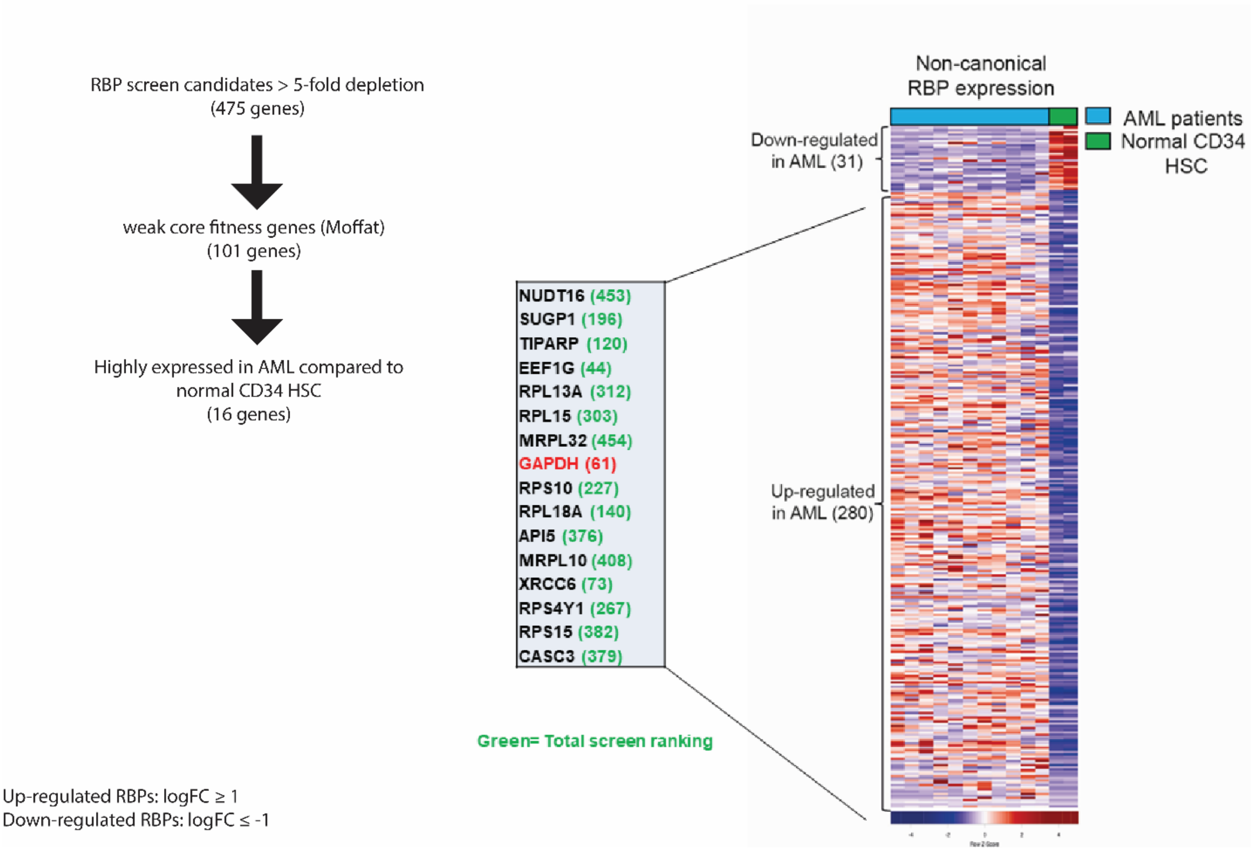


**Supplementary Figure S1: Small scale transcriptome analysis of AML patients and healthy HSC samples.** The expression pattern of non-canonical RNA binding proteins is displayed. Samples from AML patients (n = 11) and normal CD34+ hematopoietic stem cells (HSC) were analyzed. The filtering process is shown in the left panel. Up-regulation was defined as logFC ≥ 1, while down-regulation of genes was defined as logFC ≤ 1. Numbers in brackets indicate the ranking in the screen.

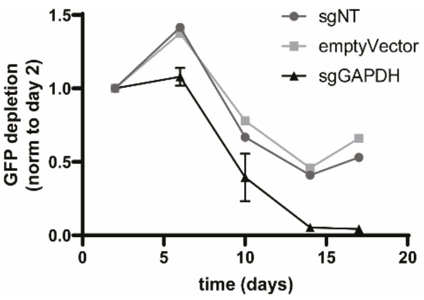


**Supplementary Figure S2: GFP competition assay of MOLM-13 cells.** Cells were transfected by an empty vector control, a sgRNA non-targeting vector and 4 different sgRNA vector targeting GAPDH (sgGAPDH). The samples treated each with one of four different sgRNAs targeting GAPDH were averaged and the SEM is displayed. GFP fluorescence was monitored over time and the fluorescence signal normalized to day 2.

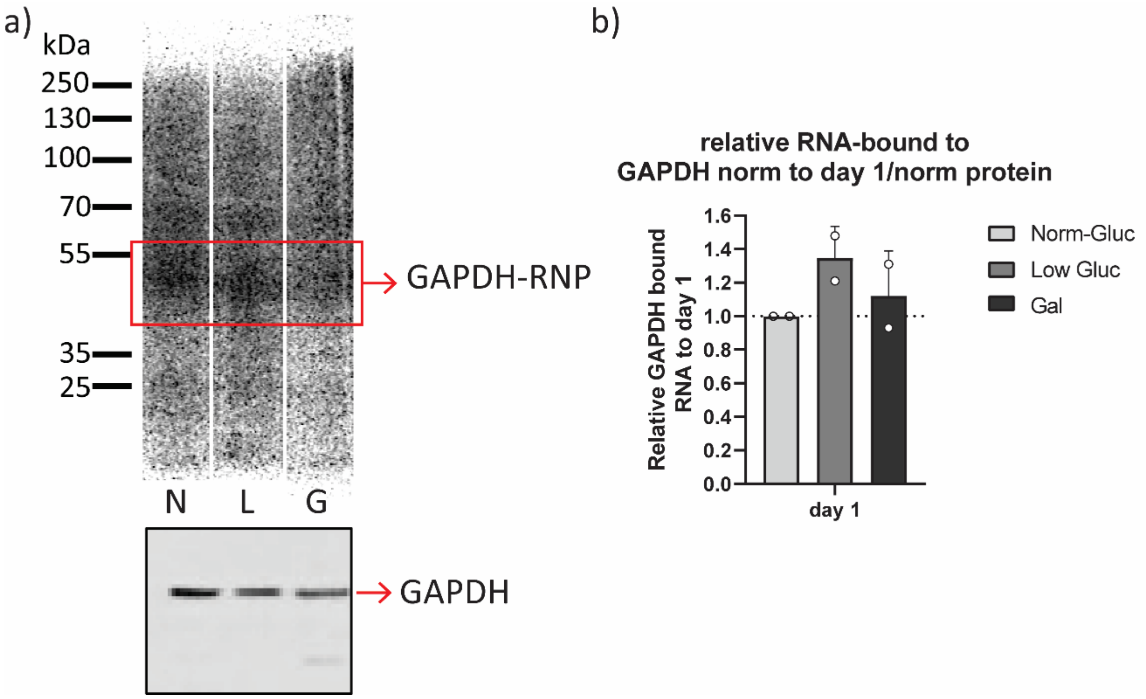


**Supplementary Figure S3: Autoradiography of GAPDH bound to RNAs.** MOLM-13 cells were cultivated in medium containing either galactose (Gal; G), normal glucose (Norm-Gluc; N) or low glucose (Low-Gluc, L) concentrations. GAPDH in complex with its bound RNA was isolated with an adapted eCLIP protocol using a polyclonal anti-GAPDH antibody **(a)** The $^{32}$P isotope RNA labeling was utilized to detect the GAPDH-RNA interaction by autoradiography. **(b)** The band intensities of the autoradiogram were quantified by densitometry. The data (n = 2) are expressed as mean ± SD.

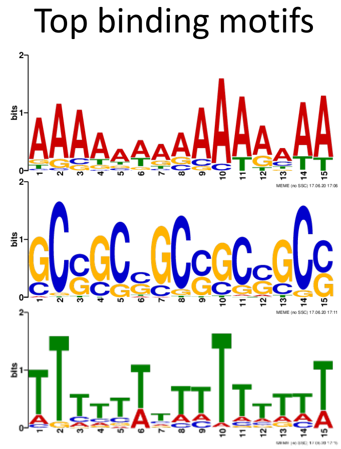


| E-value | Sites | Width |
|---|---|---|
| 1.1e-111 | 498 | 15 |
| 6.6e-128 | 871 | 15 |
| 2.5e-065 | 620 | 15 |

**Supplementary Figure S4: Identified top binding motifs based on CLIP.** Cross linking RNA immunoprecipitation (CLIP) was performed to identify GAPDH targets. The data was analyzed for the discovery of enriched binding motifs. Among the identified motifs, the three top binding motifs are displayed with their corresponding p-value. The binding motifs are exemplary for normal glucose conditions and are consistent with the conditions at low glucose concentration and in the presence of galactose.

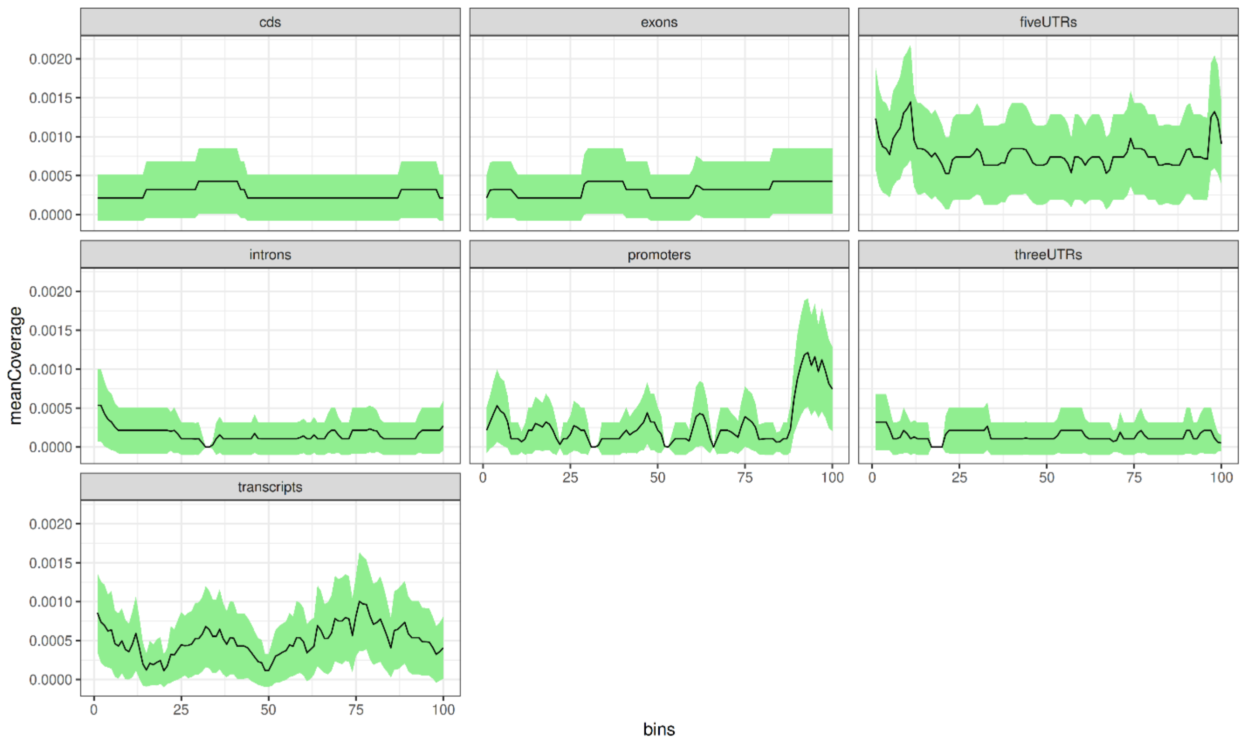


**Supplementary Figure S5: CLIP data in normal glucose conditions.** Mean coverage of CLIP peaks according to genomic localizations in normal glucose conditions.

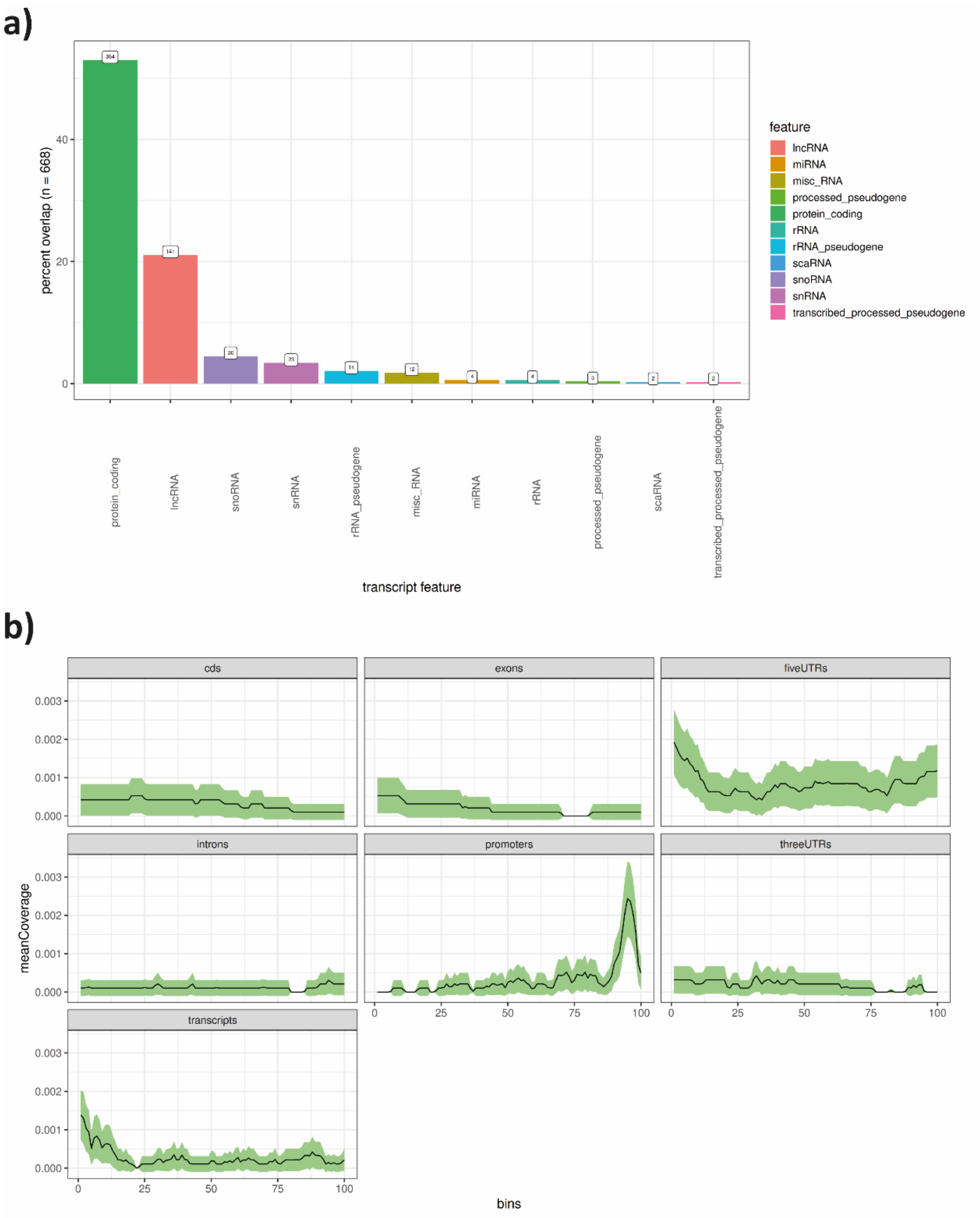


**Supplementary Figure S6: CLIP data in low glucose conditions. (a)** Genomic distribution of CLIP targets in low glucose conditions. **(b)** Mean coverage of CLIP peaks according to genomic localizations in low glucose conditions.

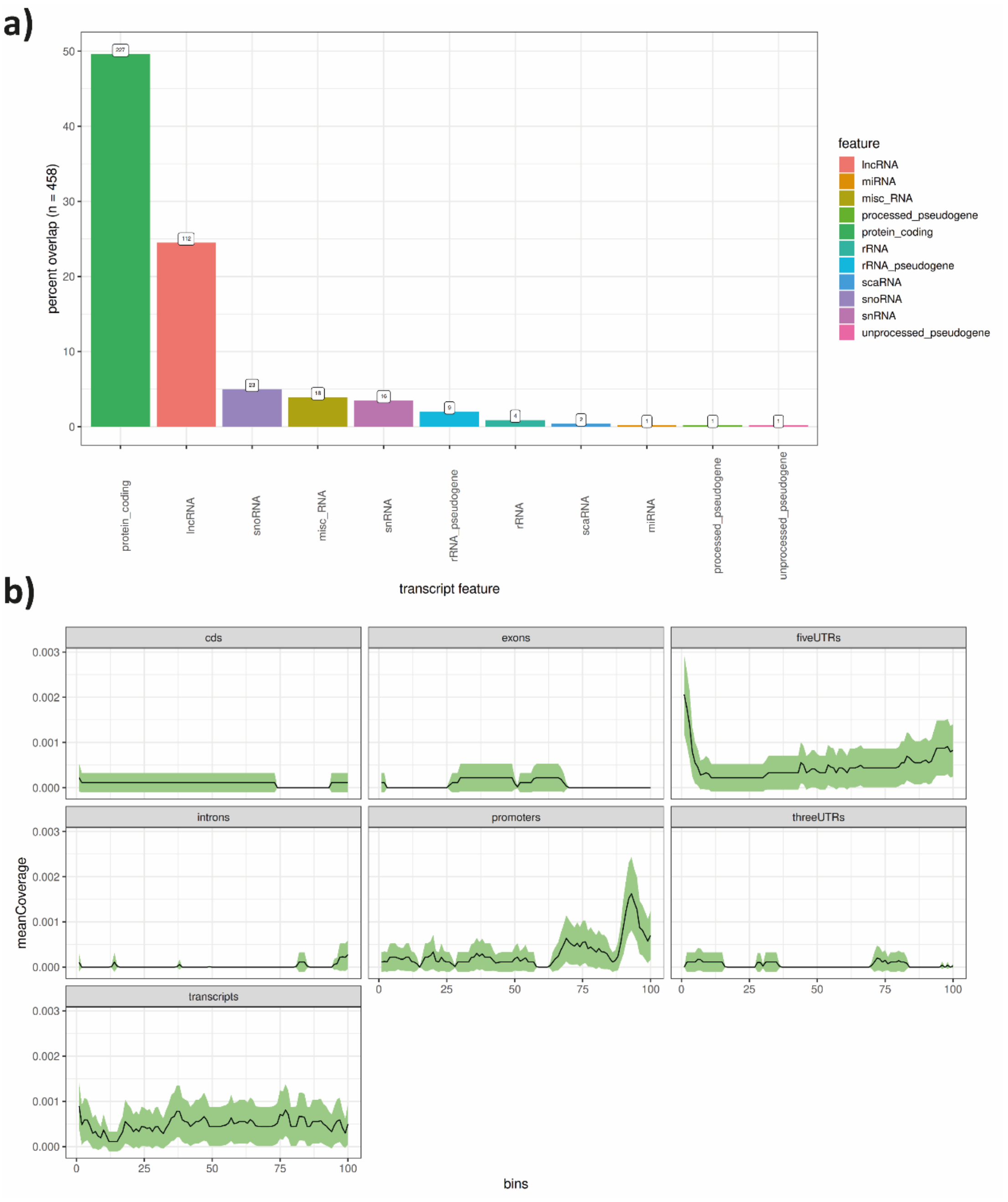


**Supplementary Figure S7: CLIP data in galactose conditions. (a)** Genomic distribution of CLIP targets in galactose conditions. **(b)** Mean coverage of CLIP peaks according to genomic localizations in galactose conditions.

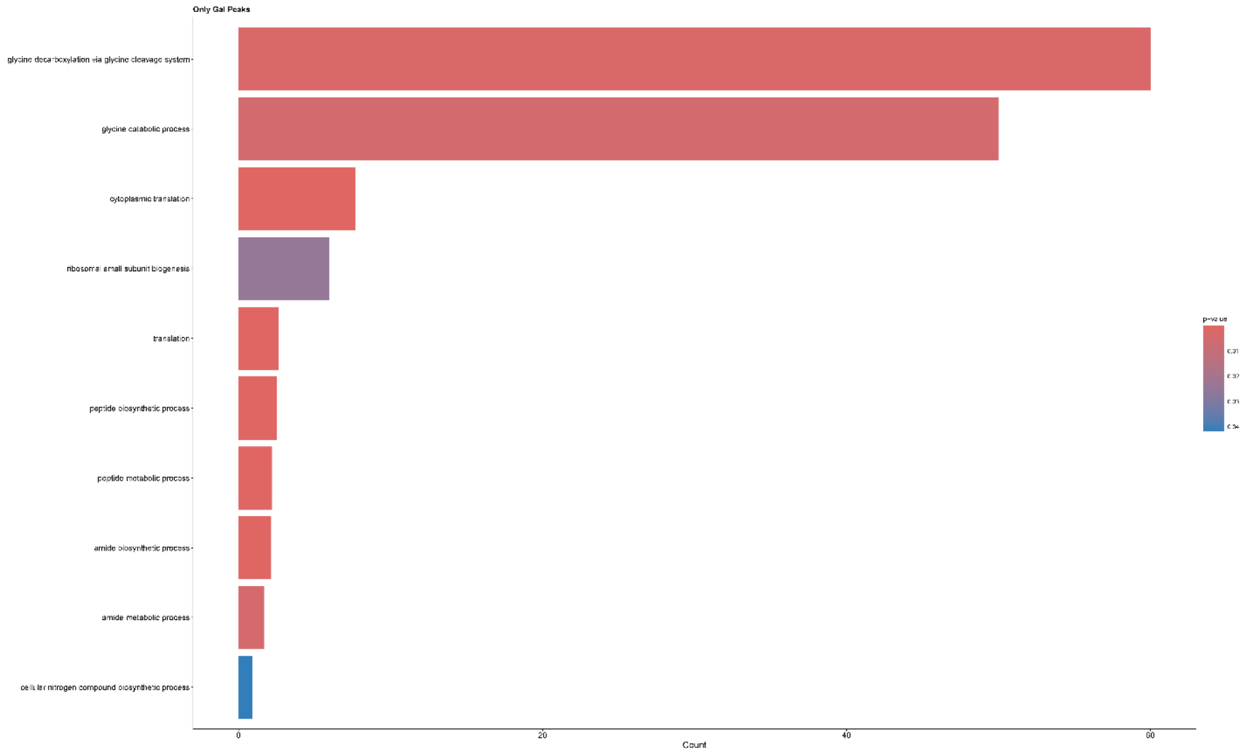


**Supplementary Figure S8: GO term analysis based on CLIP data in galactose conditions.**

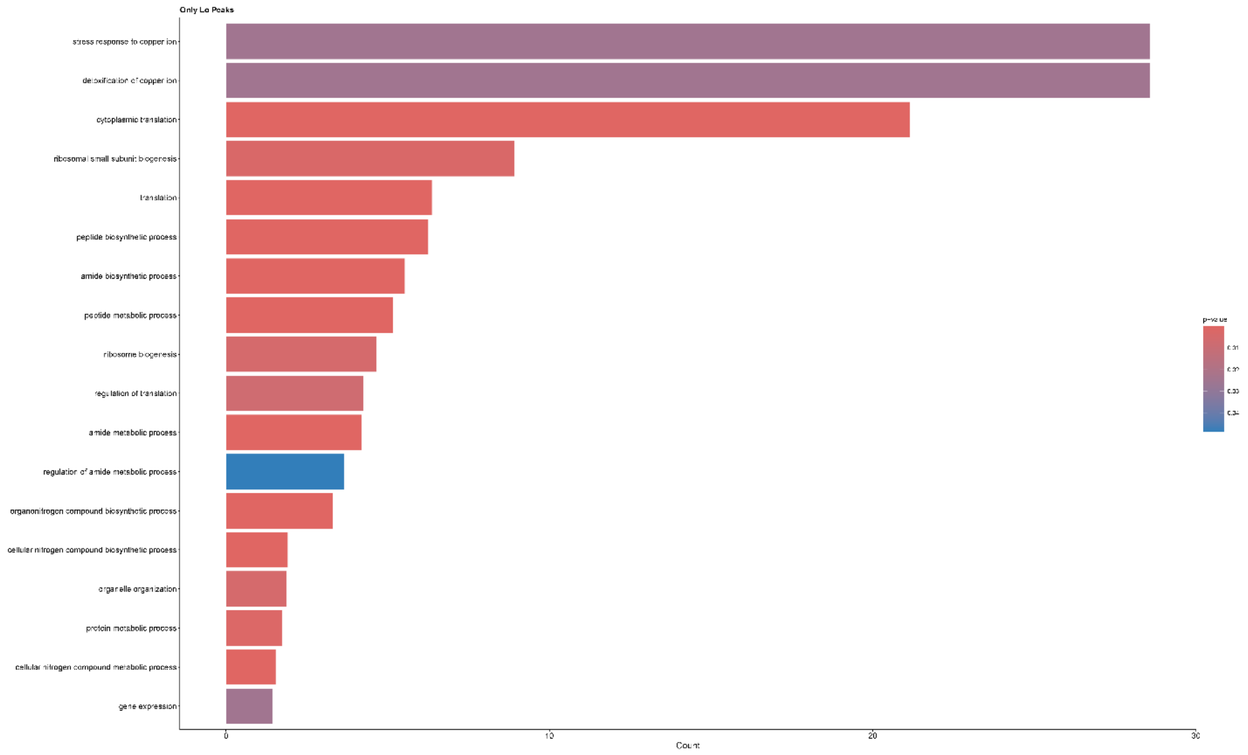


**Supplementary Figure S9: GO term analysis based on CLIP data in low glucose conditions.**

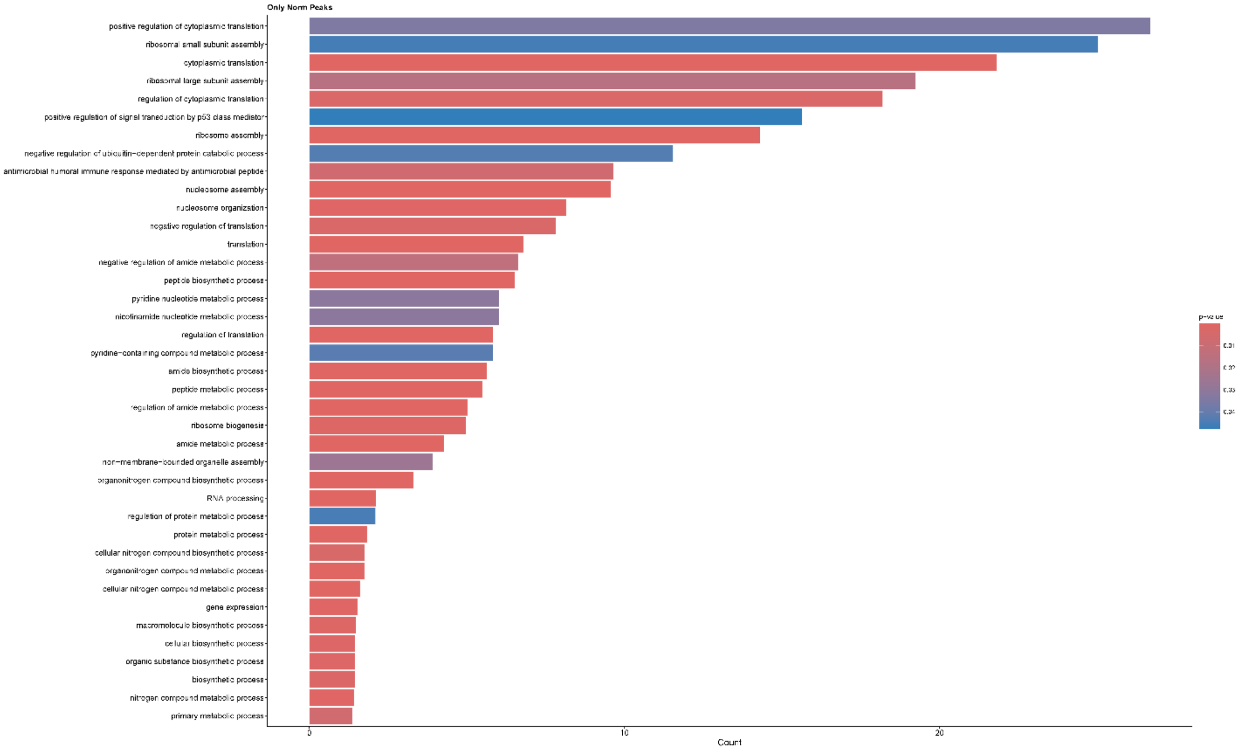


**Supplementary Figure S10: GO term analysis based on CLIP data in normal glucose conditions.**

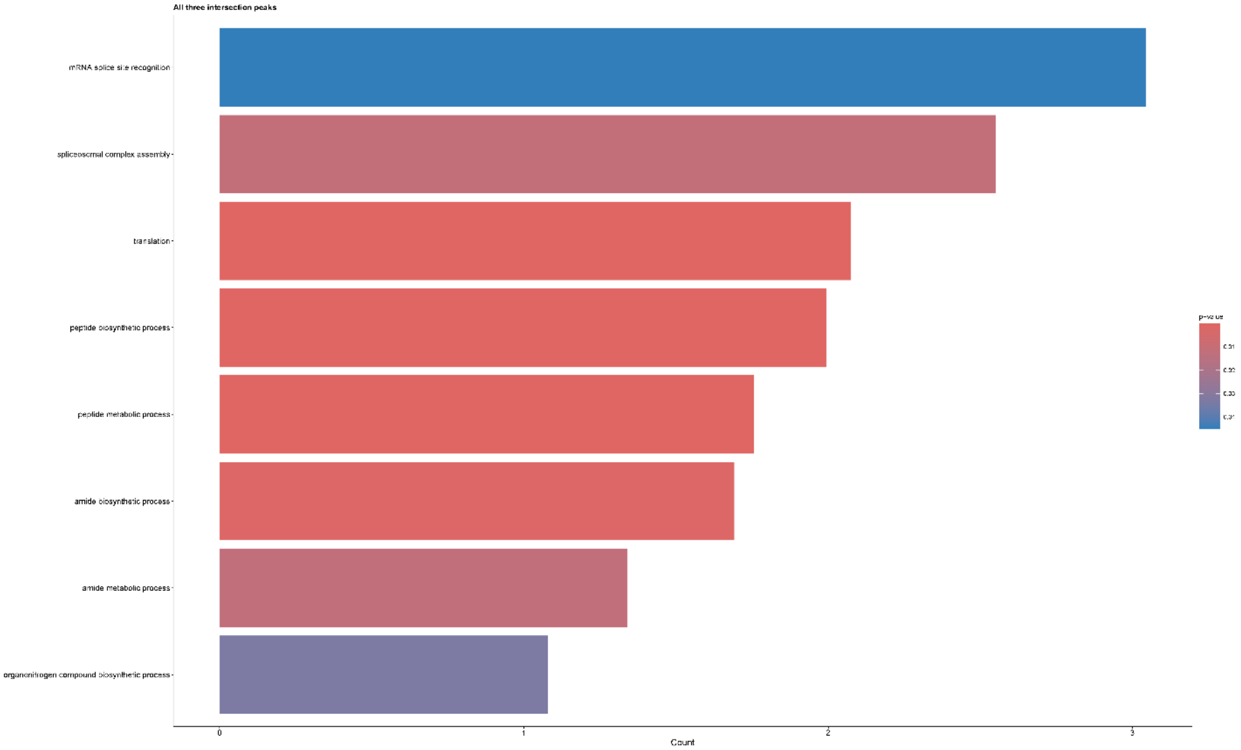


**Supplementary Figure S11: GO term analysis based on CLIP data in intersected conditions.**

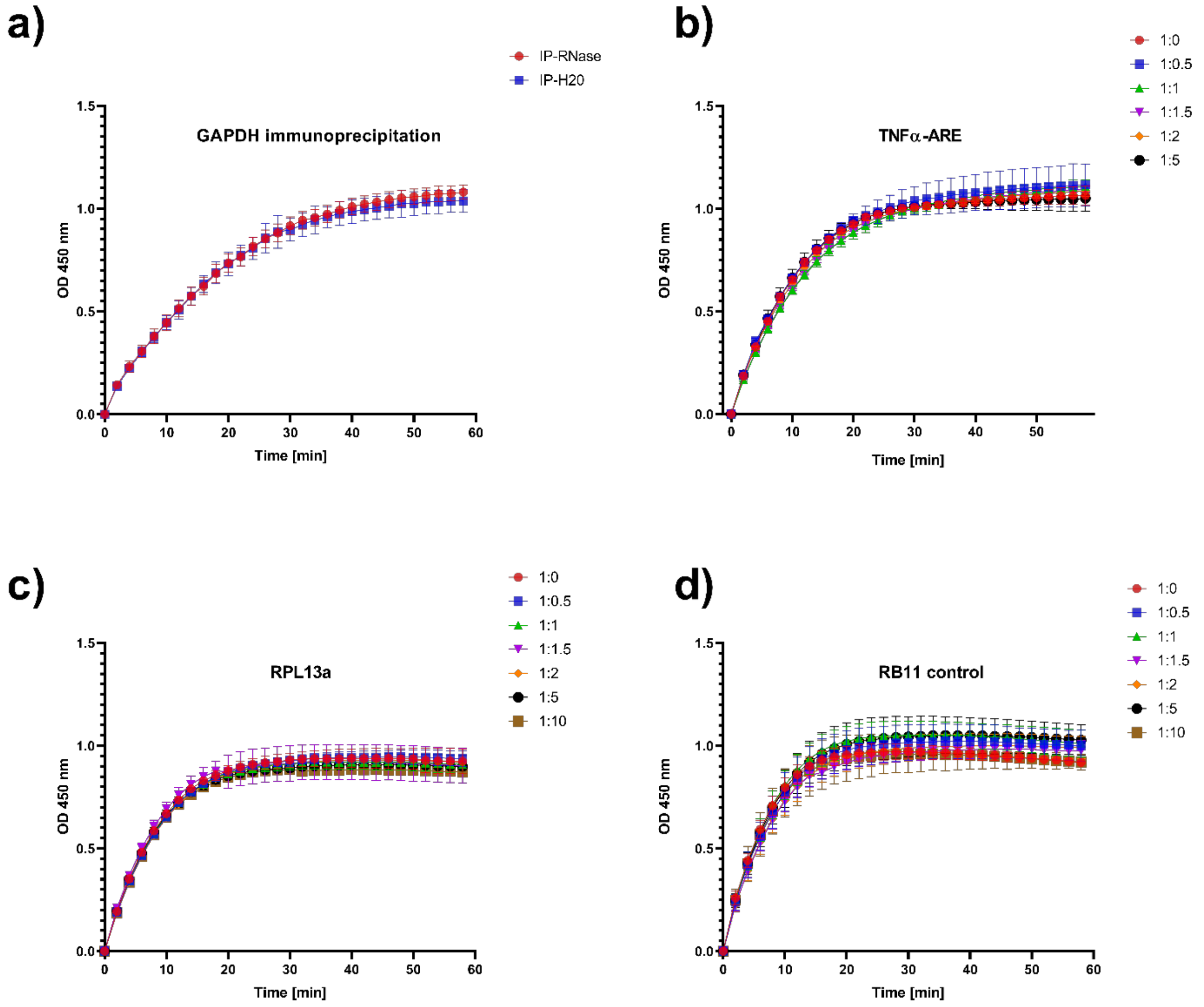


**Supplementary Figure S12: GAPDH activity assay. (a)** GAPDH immunoprecipitation (IP) was performed. GAPDH activity was detected after treating samples with or without RNase treatment. **(b)** *in vitro* synthesized TNFα-ARE was incubated with increasing concentrations of recombinant GAPDH and a GAPDH activity assay was performed. **(c)** GAPDH activity assay with *in vitro* synthesized RPL13a. **(d)** GAPDH activity assay with *in vitro* synthesized Rβ31 control.

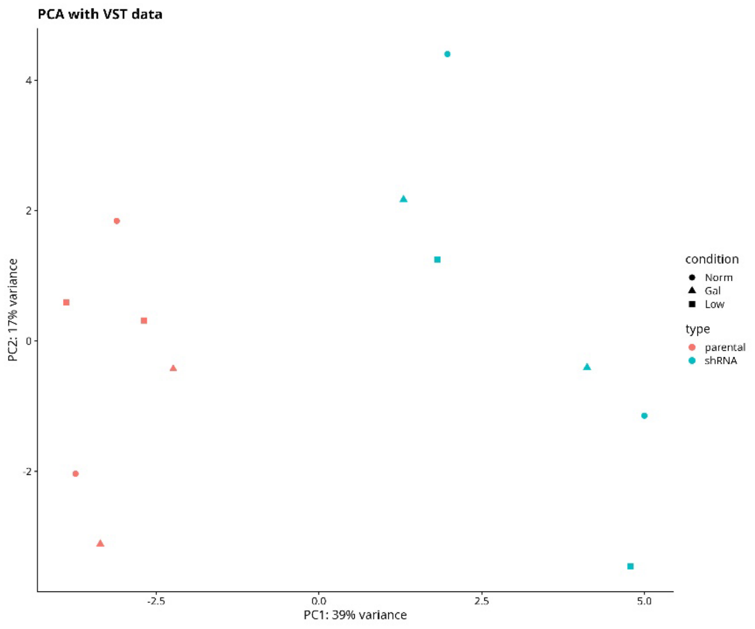


**Supplementary Figure S13: PCA plots for different metabolic conditions of RNA-knockdown RNA-sequencing.** The samples for the normal glucose (Norm), low glucose (Low) and galactose (Gal) conditions were clustered together based on the knockdown of GAPDH.

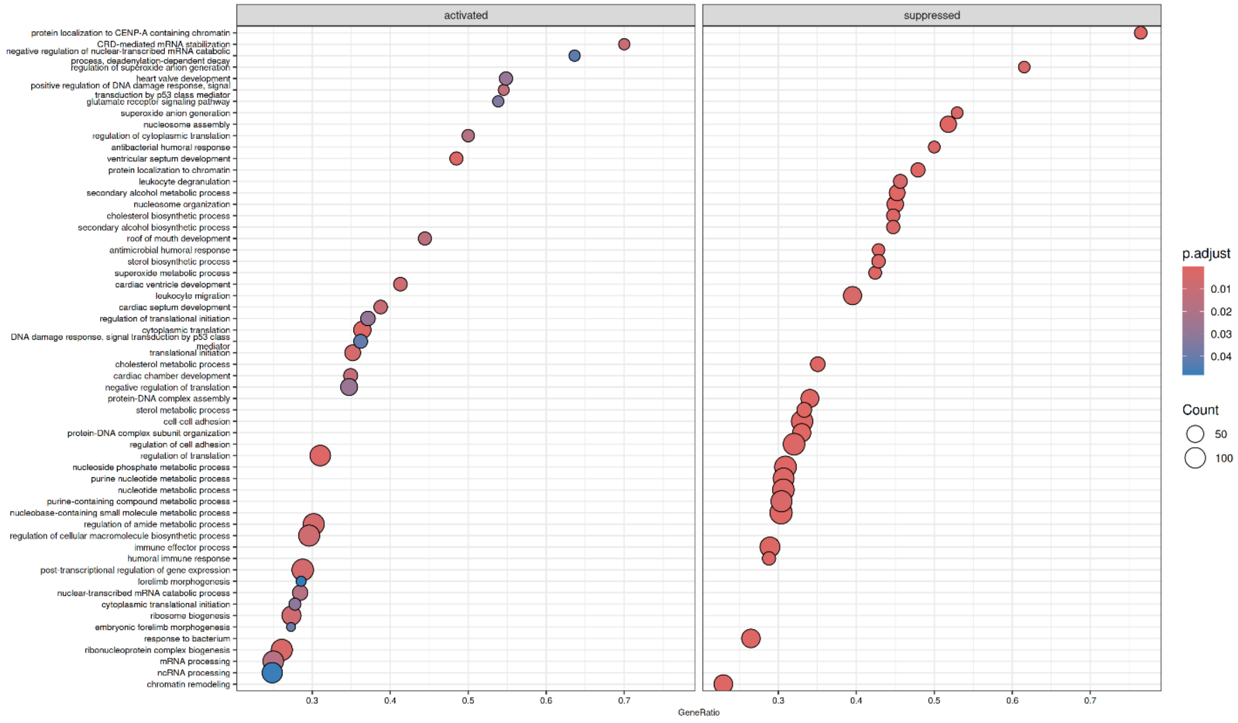


**Supplementary Figure S14: Gene set enrichment analysis after GAPDH knockdown.** The knockdown of GAPDH was realized by using shRNA targeting GAPDH in MOLM-13 cells. After RNA-seq gene set enrichment analysis was performed. Displayed are activated and suppressed gene sets.

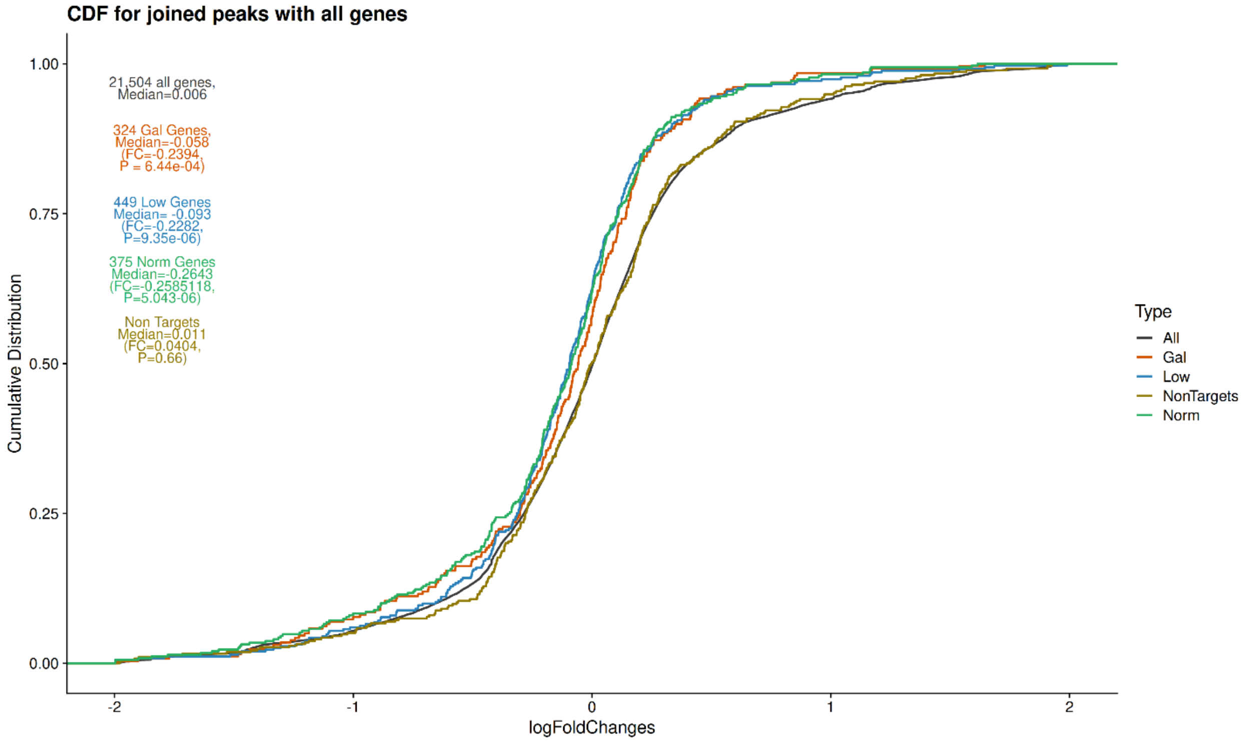


**Supplementary Figure S15: Cumulative distribution function (CDF) plot for joined peaks for all conditions.** The CDF plot for joined peaks under normal (Norm) glucose, low glucose conditions, the presence of galactose (Gal) and for all genes is shown. Bindings sites based on CLIP data were intersected with the RNA seq after GAPDH knockdown.

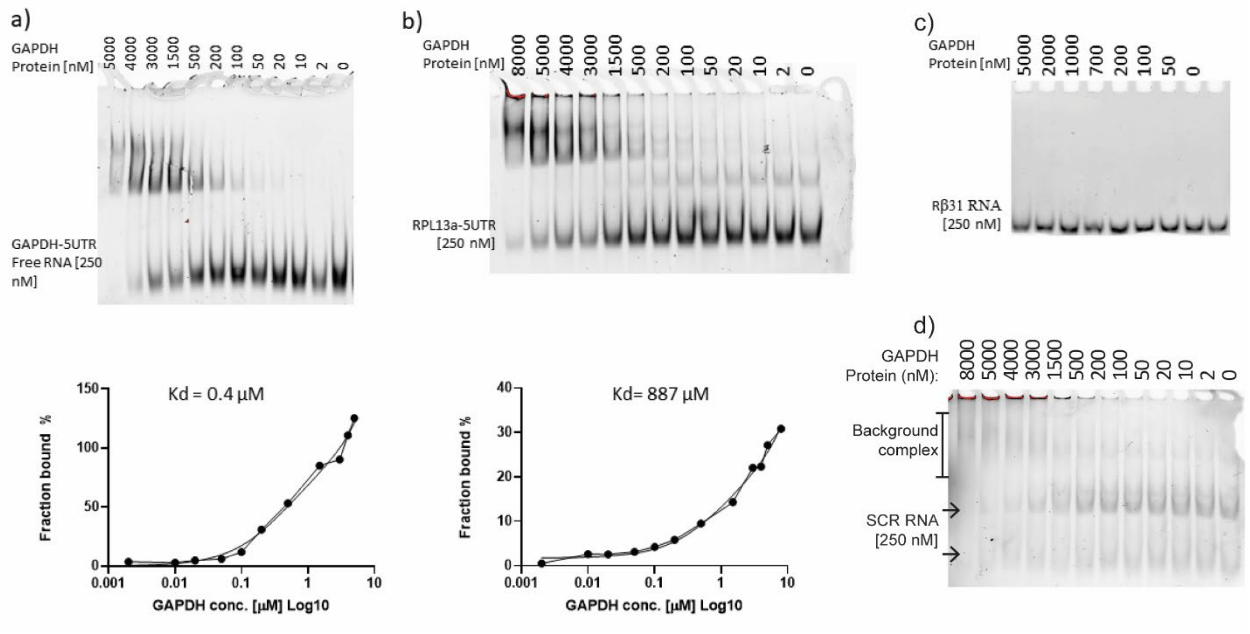


**Supplementary Figure S16: Electrophoresis mobility shift assay (EMSA) for identified CLIP targets.** Varying concentrations of recombinant produced GAPDH were with RNA targets, identified by CLIP. FITC-labeled RNA was used to visualize GAPDH-RNA interactions by EMSA. Bound fractions were calculated based on the fluorescence signal intensity. **(a)** 5'UTR of GAPDH. **(b)** 5'UTR of RPL13a. **(c)** Rβ31 control RNA. **(d)** Scrambled (SCR) control RNA.

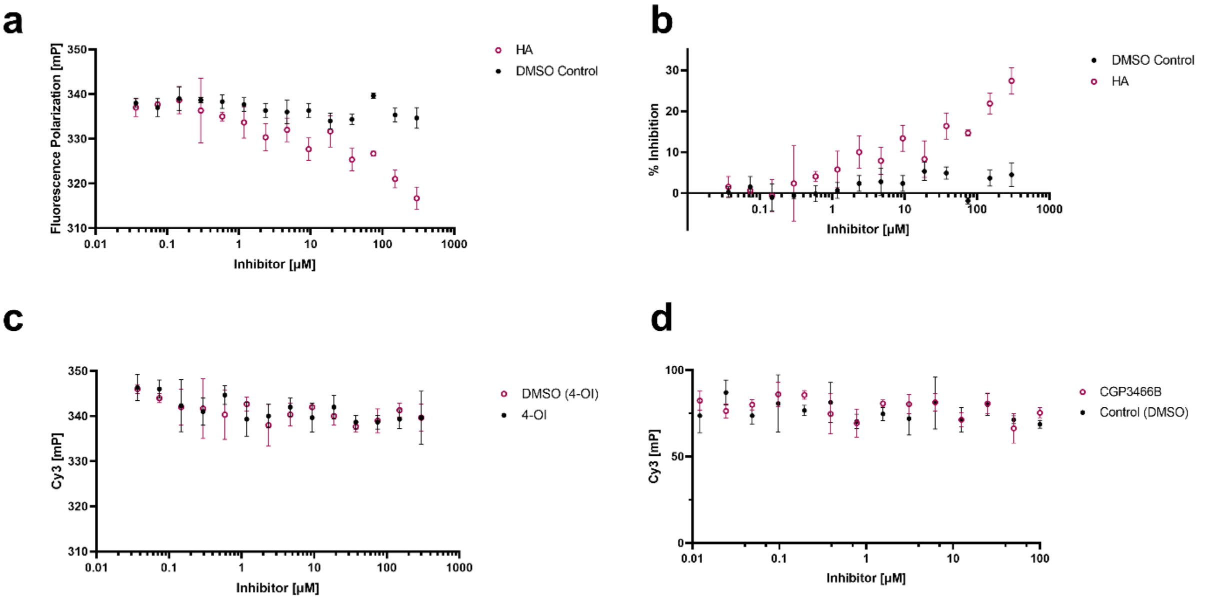


**Supplementary Figure S17: Fluorescence polarization assay. (a)** FP assay of Cy3-labeled TNFα-ARE and Rβ31 control. **(b)** Inhibitory effect expressed as percentage of inhibition. **(c)** FP assay of Cy3-labeled TNFα-ARE in the presence or absence of the GAPDH inhibitor 4-octyl itaconate (4-OI). **(d)** FP assay of Cy3-labeled TNFα-ARE in the presence or absence of the GAPDH nitrosylation blocker CGP3466B. Data are shown as mean ± SD.

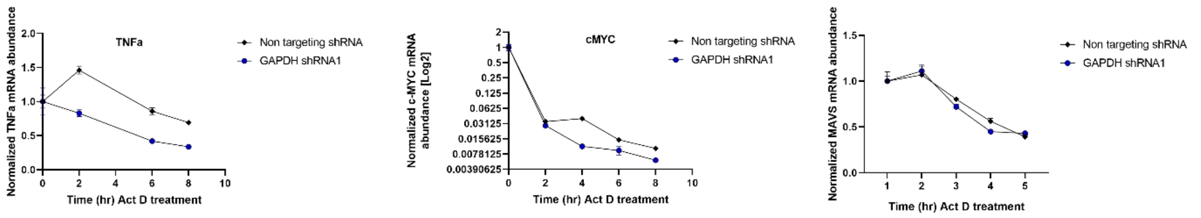


**Supplementary Figure S18: Actinomycin D (Act D) stability assay of TNFα, MYC and MAVS transcripts.** MOLM-13 cells with a GAPDH knockdown and MOLM-13 cells with a non-targeting control were treated with Act D. Expression of TNFα, MYC and MAVS transcripts was monitored over time by quantitative PCR. MAVS served as a negative control.

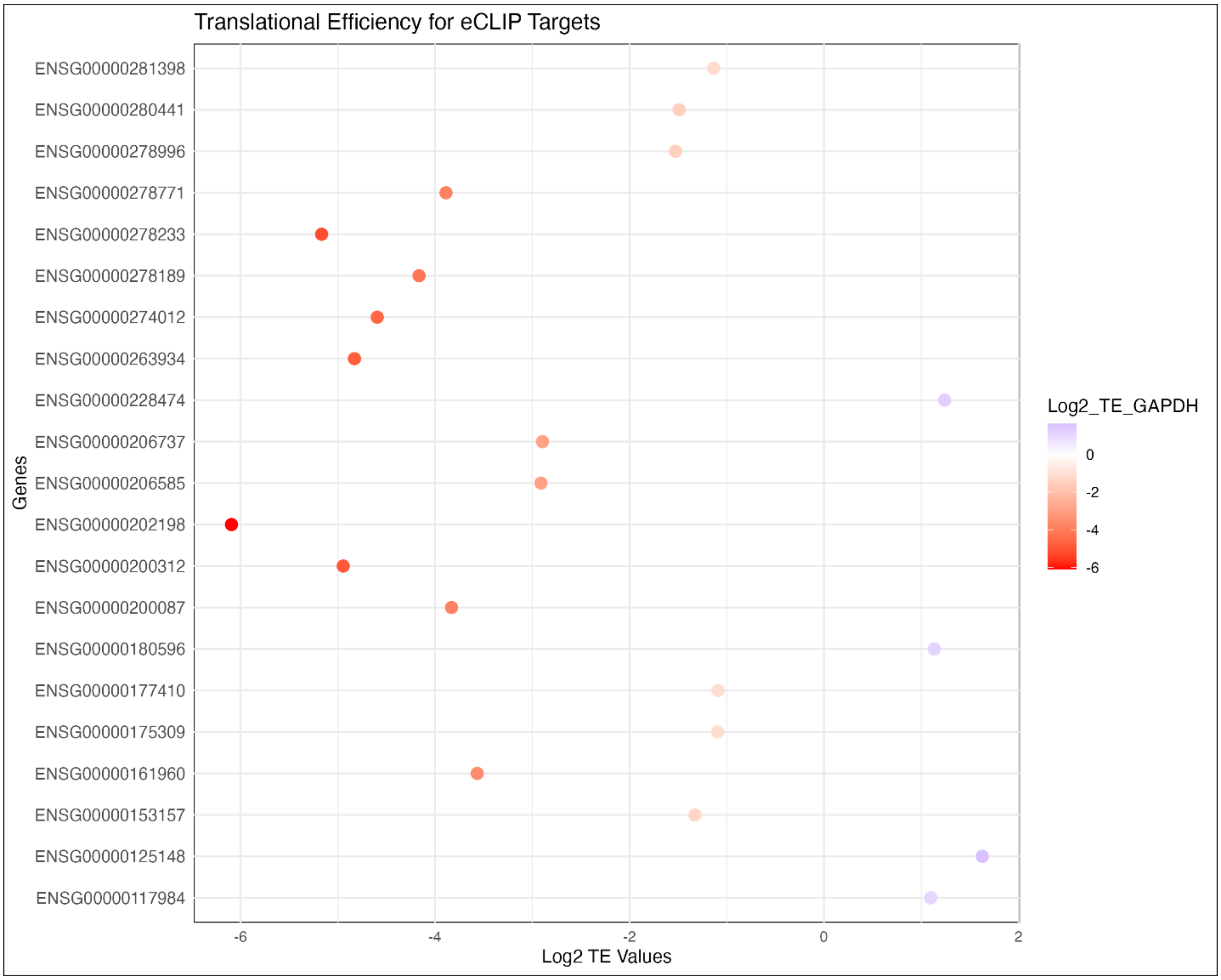


**Supplementary Figure S19: Translation efficiency of eCLIP taregts.** Based on the RNAseq data of the translation efficiency assay, genes with a TE value above +1 and below -1 are shown.

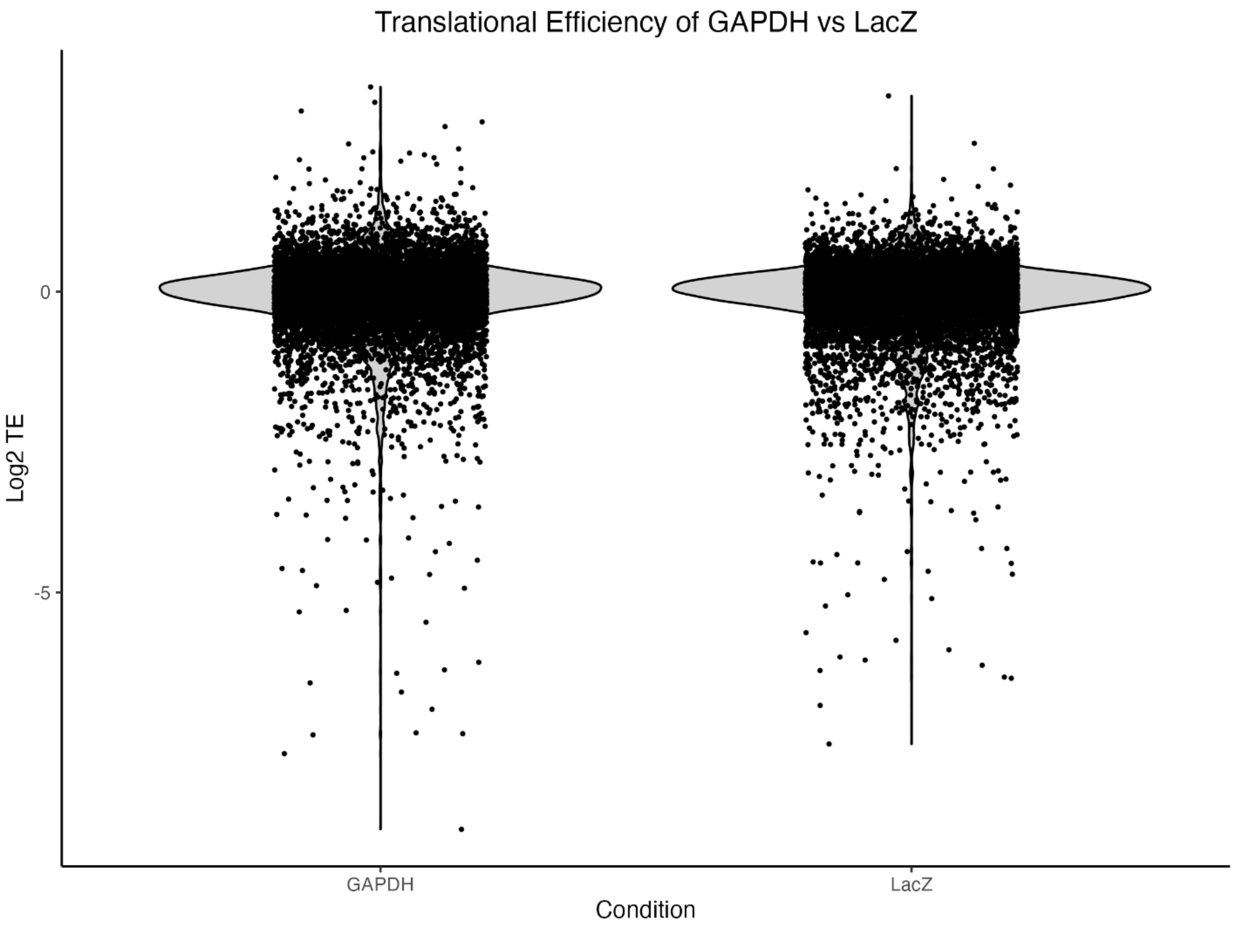


**Supplementary Figure S20: Translation efficiency of all genes.** Based on the RNAseq data of the translation efficiency assay.

**Supplementary Table S1: Raw RNA-seq reads from healthy control cells**

| Name | Platform | SRP ID | Cell Type | GSE Number |
|---|---|---|---|---|
| SRR11164713_GSM4333128_PID730_CMP_MEP | Illumina NovaSeq 6000 | SRP250479 | common myeloid/megakaryocyte-erythrocyte progenitors | GSE145802 |
| SRR11164714_GSM4333129_PID730_HSC | Illumina NovaSeq 6000 | SRP250479 | hematopoietic stem/multipotent progenitor cells | GSE145802 |
| SRR11164715_GSM4333130_PID730_CMP | Illumina NovaSeq 6000 | SRP250479 | common myeloid progenitors | GSE145802 |
| SRR11164716_GSM4333131_PID730_MEP | Illumina NovaSeq 6000 | SRP250479 | megakaryocyte-erythrocyte progenitors | GSE145802 |
| SRR11164717_GSM4333132_PID730_GMP | Illumina NovaSeq 6000 | SRP250479 | granulocyte-macrophage progenitors | GSE145802 |
| SRR11164731_GSM4333146_PID94_CMP_MEP | Illumina NovaSeq 6000 | SRP250479 | common myeloid/megakaryocyte-erythrocyte progenitors | GSE145802 |
| SRR11164732_GSM4333147_PID94_HSC | Illumina NovaSeq 6000 | SRP250479 | hematopoietic stem/multipotent progenitor cells | GSE145802 |
| SRR11164733_GSM4333148_PID94_GMP | Illumina NovaSeq 6000 | SRP250479 | granulocyte-macrophage progenitors | GSE145802 |
| SRR11164734_GSM4333149_PID757_CMP_MEP | Illumina NovaSeq 6000 | SRP250479 | common myeloid/megakaryocyte-erythrocyte progenitors | GSE145802 |
| SRR11164735_GSM4333150_PID757_GMP | Illumina NovaSeq 6000 | SRP250479 | granulocyte-macrophage progenitors | GSE145802 |
| SRR11164750_GSM4333165_Haemo_CMP_MEP | Illumina NovaSeq 6000 | SRP250479 | common myeloid/megakaryocyte-erythrocyte progenitors | GSE145802 |
| SRR11164751_GSM4333166_Haemo_HSC | Illumina NovaSeq 6000 | SRP250479 | hematopoietic stem/multipotent progenitor cells | GSE145802 |

| | | | | |
|---|---|---|---|---|
| SRR11164752_GSM4333167_Haemo_CMP | Illumina NovaSeq 6000 | SRP250479 | common myeloid progenitors | GSE145802 |
| SRR11164753_GSM4333168_Haemo_MEP | Illumina NovaSeq 6000 | SRP250479 | megakaryocyte-erythrocyte progenitors | GSE145802 |
| SRR11164754_GSM4333169_Haemo_GMP | Illumina NovaSeq 6000 | SRP250479 | granulocyte-macrophage progenitors | GSE145802 |
| SRR11164760_GSM4333175_PID781_CMP_MEP | Illumina NovaSeq 6000 | SRP250479 | common myeloid/megakaryocyte-erythrocyte progenitors | GSE145802 |
| SRR11164761_GSM4333176_PID781_HSC | Illumina NovaSeq 6000 | SRP250479 | hematopoietic stem/multipotent progenitor cells | GSE145802 |
| SRR11164762_GSM4333177_PID781_CMP | Illumina NovaSeq 6000 | SRP250479 | common myeloid progenitors | GSE145802 |
| SRR11164763_GSM4333178_PID781_MEP | Illumina NovaSeq 6000 | SRP250479 | megakaryocyte-erythrocyte progenitors | GSE145802 |
| SRR11164764_GSM4333179_PID781_GMP | Illumina NovaSeq 6000 | SRP250479 | granulocyte-macrophage progenitors | GSE145802 |
| SRR11164798_GSM4333213_BC1_CMP_MEP | Illumina NovaSeq 6000 | SRP250479 | common myeloid/megakaryocyte-erythrocyte progenitors | GSE145802 |
| SRR11164799_GSM4333214_BC1_HSC | Illumina NovaSeq 6000 | SRP250479 | hematopoietic stem/multipotent progenitor cells | GSE145802 |
| SRR11164800_GSM4333215_BC1_CMP | Illumina NovaSeq 6000 | SRP250479 | common myeloid progenitors | GSE145802 |
| SRR11164801_GSM4333216_BC1_MEP | Illumina NovaSeq 6000 | SRP250479 | megakaryocyte-erythrocyte progenitors | GSE145802 |
| SRR11164802_GSM4333217_BC1_GMP | Illumina NovaSeq 6000 | SRP250479 | granulocyte-macrophage progenitors | GSE145802 |

| | | | | |
|---|---|---|---|---|
| SRR11164803_GSM4333218_BC2_CMP_MEP | Illumina NovaSeq 6000 | SRP250479 | common myeloid/megakaryocyte-erythrocyte progenitors | GSE145802 |
| SRR11164804_GSM4333219_BC2_HSC | Illumina NovaSeq 6000 | SRP250479 | hematopoietic stem/multipotent progenitor cells | GSE145802 |
| SRR11164805_GSM4333220_BC2_CMP | Illumina NovaSeq 6000 | SRP250479 | common myeloid progenitors | GSE145802 |
| SRR11164806_GSM4333221_BC2_MEP | Illumina NovaSeq 6000 | SRP250479 | megakaryocyte-erythrocyte progenitors | GSE145802 |
| SRR11164807_GSM4333222_BC2_GMP | Illumina NovaSeq 6000 | SRP250479 | granulocyte-macrophage progenitors | GSE145802 |
| SRR11164808_GSM4333223_BC3_CMP_MEP | Illumina NovaSeq 6000 | SRP250479 | common myeloid/megakaryocyte-erythrocyte progenitors | GSE145802 |
| SRR11164809_GSM4333224_BC3_HSC | Illumina NovaSeq 6000 | SRP250479 | hematopoietic stem/multipotent progenitor cells | GSE145802 |
| SRR11601147_GSM4403509_HSPC_1_RNA-seq | Illumina HiSeq 4000 | SRP258171 | Hematopoietic stem and progenitor cells (HSPCs) | GSE149237 |
| SRR11601148_GSM4403509_HSPC_1_RNA-seq | Illumina HiSeq 4000 | SRP258171 | Hematopoietic stem and progenitor cells (HSPCs) | GSE149237 |
| SRR11601149_GSM4403510_HSPC_2_RNA_seq | Illumina HiSeq 4000 | SRP258171 | Hematopoietic stem and progenitor cells (HSPCs) | GSE149237 |
| SRR11601150_GSM4403510_HSPC_2_RNA_seq | Illumina HiSeq 4000 | SRP258171 | Hematopoietic stem and progenitor cells (HSPCs) | GSE149237 |
| SRR11601151_GSM4403511_HSPC_3_RNA-seq | Illumina HiSeq 4000 | SRP258171 | Hematopoietic stem and progenitor cells (HSPCs) | GSE149237 |
| SRR11601152_GSM4403511_HSPC_3_RNA-seq | Illumina HiSeq 4000 | SRP258171 | Hematopoietic stem and progenitor cells (HSPCs) | GSE149237 |
| SRR11601153_GSM4403512_HSPC_4_RNA_seq | Illumina HiSeq 4000 | SRP258171 | Hematopoietic stem and progenitor cells (HSPCs) | GSE149237 |
| SRR11601154_GSM4403512_HSPC_4_RNA_seq | Illumina HiSeq 4000 | SRP258171 | Hematopoietic stem and progenitor cells (HSPCs) | GSE149237 |
| SRR11601155_GSM4403513_HSPC_5_RNA-seq | Illumina HiSeq 4000 | SRP258171 | Hematopoietic stem and progenitor cells (HSPCs) | GSE149237 |

| | | | | |
|---|---|---|---|---|
| SRR11601156_GSM4403513_HSPC_5_RNA-seq | Illumina HiSeq 4000 | SRP258171 | Hematopoietic stem and progenitor cells (HSPCs) | GSE149237 |
| SRR14301187_GSM5259993_library7 | Illumina NovaSeq 6000 | SRP315886 | CD34+ hematopoietic stem/progenitor cells (HSPCs) | GSE149237 |
| SRR14301188_GSM5259994_library8 | Illumina NovaSeq 6000 | SRP315886 | CD34+ hematopoietic stem/progenitor cells (HSPCs) | GSE149237 |
| SRR14301189_GSM5259995_library9 | Illumina NovaSeq 6000 | SRP315886 | CD34+ hematopoietic stem/progenitor cells (HSPCs) | GSE149237 |
| SRR6464417_GSM2931519_BM_HSC_1 | Illumina HiSeq 2000 | SRP128918 | Bone marrow hematopoietic stem cells | GSE109093 |
| SRR6464418_GSM2931520_BM_HSC_2 | Illumina HiSeq 2000 | SRP128918 | Bone marrow hematopoietic stem cells | GSE109093 |
| SRR6464419_GSM2931521_BM_HSC_3 | Illumina HiSeq 2000 | SRP128918 | Bone marrow hematopoietic stem cells | GSE109093 |
| SRR6464426_GSM2931528_BM_PROG_1 | Illumina HiSeq 2000 | SRP128918 | Bone marrow hematopoietic progenitor cells | GSE109093 |
| SRR6464427_GSM2931529_BM_PROG_2 | Illumina HiSeq 2000 | SRP128918 | Bone marrow hematopoietic progenitor cells | GSE109093 |
| SRR6464428_GSM2931530_BM_PROG_3 | Illumina HiSeq 2000 | SRP128918 | Bone marrow hematopoietic progenitor cells | GSE109093 |
| SRR2753085 | NextSeq 500 | SRP065216 | lymphoid-primed multipotent progenitor cell | GSE75384 |
| SRR2753090 | NextSeq 500 | SRP065216 | common myeloid progenitor cell | GSE75384 |
| SRR2753091 | NextSeq 500 | SRP065216 | granulocyte macrophage progenitor cell | GSE75384 |
| SRR2753092 | NextSeq 500 | SRP065216 | hematopoietic stem cell | GSE75384 |
| SRR2753093 | NextSeq 500 | SRP065216 | megakaryocyte erythroid progenitor cell | GSE75384 |
| SRR2753095 | NextSeq 500 | SRP065216 | multipotent progenitor cell | GSE75384 |
| SRR2753096 | NextSeq 500 | SRP065216 | common myeloid progenitor cell | GSE75384 |
| SRR2753097 | NextSeq 500 | SRP065216 | granulocyte macrophage progenitor cell | GSE75384 |
| SRR2753098 | NextSeq 500 | SRP065216 | hematopoietic stem cell | GSE75384 |

| | | | | |
|---|---|---|---|---|
| SRR2753099 | NextSeq 500 | SRP065216 | megakaryocyte erythroid progenitor cell | GSE75384 |
| SRR2753101 | NextSeq 500 | SRP065216 | multipotent progenitor cell | GSE75384 |
| SRR2753104 | NextSeq 500 | SRP065216 | common myeloid progenitor cell | GSE75384 |
| SRR2753105 | NextSeq 500 | SRP065216 | granulocyte macrophage progenitor cell | GSE75384 |
| SRR2753106 | NextSeq 500 | SRP065216 | hematopoietic stem cell | GSE75384 |
| SRR2753107 | NextSeq 500 | SRP065216 | lymphoid-primed multipotent progenitor cell | GSE75384 |
| SRR2753108 | NextSeq 500 | SRP065216 | megakaryocyte erythroid progenitor cell | GSE75384 |
| SRR2753110 | NextSeq 500 | SRP065216 | multipotent progenitor cell | GSE75384 |
| SRR2753114 | NextSeq 500 | SRP065216 | common myeloid progenitor cell | GSE75384 |
| SRR2753115 | NextSeq 500 | SRP065216 | granulocyte macrophage progenitor cell | GSE75384 |
| SRR2753116 | NextSeq 500 | SRP065216 | hematopoietic stem cell | GSE75384 |
| SRR2753117 | NextSeq 500 | SRP065216 | lymphoid-primed multipotent progenitor cell | GSE75384 |
| SRR2753118 | NextSeq 500 | SRP065216 | megakaryocyte erythroid progenitor cell | GSE75384 |
| SRR2753120 | NextSeq 500 | SRP065216 | multipotent progenitor cell | GSE75384 |

**Supplementary Table S2: Oligo list**

| Oligos | Sequence |
|---|---|
| Array_Fwd | ttatatatcttgtggaaaggacgaaacaccg |
| Array_Rev | agccttattttaacttgctatttctagctctaaaac |
| GAPDH_F | agccacatcgctcagacac |
| GAPDH_R | gcccaatacgaccaaatct |
| TNF-FW | cagcctcttctccttcctgat |
| TNF-RE | gccagagggctgattagaga |
| MYC-FW | tgctccatgaggagacacc |
| MYC-RE | cttttccacagaaacaacatcg |
| RPL13a-FW | ggtcgtacgctgtgaaggcatc |
| RPL13a-RV | tgcggaggaaagccaggtactt |
| PKM-FW | tcctcaccaagtctggcaggtc |
| PKM-RV | gggattccgggtcacagcaatg |
| 18S-F | acccgttgaaccccattcgtga |
| 18S-R | gcctcactaaaccatccaatcgg |
| beta actin-FW | ccaaccgcgagaagatga |
| beta actin-RW | ccagaggcgtacagggatag |
| TNFα-ARE38 | gugauuauuuauuauuuauuuauuauuuauuuauuuag |
| Rβ31 | uggccaaugcccuggcucacaaauaccacug |

**Supplementary Table S3: Templates for *in vitro* transcription**

| | |
|---|---|
| GAPDH 5-UTR | taatacgactcactatagggctctctgctcctcctgttcgacagtcagccgcatcttcttttgcgtcgccagccgagccacatcgctcagacacc |
| RPL13a | taatacgactcactatagggcctccttttccaagcggctgccgaagatggcggaggtgcaggtcctggtgcttgatggtcgaggccatctcctgggccgcctggcggccatcgtggctaaacag |

**Supplementary Table S5: gRNA list**

| | |
|---|---|
| GAPDH_sg4001 | tgctggcgctgagtacgtcg |
| GAPDH_sg4002 | actgtggcgtgatggccgcg |
| GAPDH_sg4003 | tcacacccatgacgaacatg |
| GAPDH_sg4004 | ctgtaggctcatttgcaggg |
